# Human CCR4-NOT globally regulates gene expression and is a novel silencer of retrotransposon activation

**DOI:** 10.1101/2024.09.10.612038

**Authors:** Shardul Kulkarni, Alexis Morrissey, Aswathy Sebastian, Belinda Giardine, Courtney Smith, Oluwasegun T. Akinniyi, Cheryl A. Keller, Alexei Arnaoutov, Istvan Albert, Shaun Mahony, Joseph C. Reese

**Author notes:** **Corresponding Author:** Joseph C. Reese, Center for Eukaryotic Gene Regulation, Biochemistry and Molecular Biology, Penn State University, 463A North Frear Laboratory, University Park, PA, 16802.

## Abstract

CCR4-NOT regulates multiple steps in gene regulation and has been well studied in budding yeast, but much less is known about the human complex. Auxin-induced degradation was used to rapidly deplete the scaffold subunit CNOT1, and CNOT4, to characterize the functions of human CCR4-NOT in gene regulation. Depleting CNOT1 increased RNA levels and caused a widespread decrease in RNA decay. In contrast, CNOT4 depletion only modestly changed steady-state RNA levels and, surprisingly, led to a global acceleration in mRNA decay. Further, depleting either subunit resulted in a global increase in RNA synthesis. In contrast to most of the genome, the transcription of KRAB-Zinc-Finger-protein (KZNFs) genes, especially those on chromosome 19, was repressed. KZNFs are transcriptional repressors of retrotransposable elements (rTEs), and consistent with the decreased KZNFs expression, rTEs, mainly Long Interspersed Nuclear Elements (LINEs), were activated. These data establish CCR4-NOT as a global regulator of gene expression and a novel silencer of rTEs.

## Introduction

CCR4-NOT is a highly conserved complex that regulates multiple steps in gene expression in the nucleus and cytoplasm. Subunits of the Ccr4-Not complex were first identified and characterized in yeast as regulators of the TATA-binding protein and transcription initiation, but it has since been shown to regulate multiple stages of gene expression ^1–3^. However, comparatively less is known about its function in metazoans, especially its nuclear functions. In the cytoplasm, CCR4-NOT functions as the major mRNA deadenylase that is the rate limiting step in mRNA decay^4–6^. CCR4-NOT associates with ribosomes, monitors the translatability of mRNAs, and directs the destruction of untranslatable messages and those enriched in non-optimal codons^1–3,6–8^. Human CCR4-NOT contains one of either CNOT7/CNOT8 DEDD type deadenylases (yCaf1) and one of either CNOT6/CNOT6L exonuclease-endonuclease-phosphatase [EEP] family of deadenylases (yCcr4)^4,9^. It is recruited to target mRNAs via RNA binding proteins (RBP) or the miRNA machinery to specific sequences within the 3’UTRs. The large scaffold subunit CNOT1 is critical to bridge the RBP and the deadenylase subunits to target mRNAs for degradation; thus, it plays a crucial role in the function and integrity of the complex^10–14^.

The yeast and mammalian Ccr4-Not complexes are overall similar, but there are some key differences, including being composed of different subunits^1–3^. One difference is that yeast Not4p is an integral subunit of the complex, but CNOT4 in metazoans does not co-purify with the complex. yNot4/hCNOT4 is a RING domain-containing E3 ligase^3,9,15^. Since CNOT4 is not tightly associated with CCR4-NOT, its precise functions within the complex are unknown. Several studies have interrogated the function of human CCR4-NOT subunits in mRNA degradation using days-long shRNA/RNAi treatments and knockout cell lines and organisms. Most studies have focused on mRNA decay functions by targeting the catalytic deadenylases CNOT6/6L and CNOT7/8^1,16–18^. An analysis of its transcription functions and that of CNOT4 in gene expression has not been undertaken.

LINEs (Long Interspersed Nuclear Elements) are abundant retrotransposons that impact a range of functions in development, genome integrity and genome evolution^19–21^. LINE-1 (L1) sequences comprise nearly 20% of the human and mouse genomes^22^ and since L1s are responsible for the amplification of other repeat elements like SINEs, L1 activity possibly accounts for as much as 50% of mammalian DNA^23^. Activation of LINE elements is associated with the genome instability, cancer, neurological problems and aging^21,24,25^. LINEs and other repeat elements are silenced by the Kruppel-associated box zinc-finger proteins (KZFPs), the largest family of transcriptional repressors^26,27^. KZFPs recruit a transcriptional repressor complex, KAP1/TRIM28, to silence gene and LINE expression by recruiting chromatin corepressors like the SETDB1 histone methyltransferase, Heterochromatin protein 1; and the NuRD complex^26,28–31^. KZFPs also bind near promoters of protein coding genes and endogenous retrovirus LTRs to repress gene expression^32–35^. Given the importance of KZFPs in shaping the transcriptional landscape of genomes and the dire consequences of activating LINE elements, identifying the mechanisms of LINE silencing and the pathways that regulate KZFP activity is crucial to understanding gene expression, development, and genome integrity.

We employed an auxin-inducible degron (AID) strategy to efficiently and rapidly deplete CNOT1 and CNOT4 to address the function of CCR4-NOT in gene expression. We found that depleting CNOT1 stabilized mRNAs in the cell, but remarkably, depleting CNOT4 had the opposite effect, accelerating mRNA turnover. Thus, CNOT1 and CNOT4 have unique functions. Surprisingly, depleting either subunit resulted in a rapid and widespread increase in transcription. Interestingly, inactivating CCR4-NOT caused the rapid downregulation of KZFP gene expression and the activation of LINE and other retrotransposon elements throughout the genome. Thus, we have discovered a novel function for CCR4-NOT in regulating genome-wide transcription and maintaining genome integrity by suppressing the activation of transposable elements.

## Results

### An Auxin inducible degron (AID) system for rapid degradation of human CNOT1 and CNOT4

CRISPR/Cas9 editing introduced an AID tag at the C-terminus of CNOT1 and CNOT4 in colorectal adenocarcinoma DLD-1^TIR1+^ cells (Supplementary Figure S1A-C), a stably pseudodiploid adherent cell line with a modal chromosome number of 46. CNOT1 is the central scaffold subunit and CNOT4 is the E3-ubiquitin ligase subunit of the complex. Cells containing AID-tagged subunits displayed normal viability and growth (Supplementary Figure S1D) and adding the AID tag did not affect the expression levels of other CCR4-NOT proteins (Supplementary Figure S1E-F) in the absence of auxin. AID-tagged proteins can display constitutive degradation without auxin, but the expression levels of the tagged proteins were comparable to those of untagged proteins (Figure 1A, 1B). Importantly, adding auxin led to a rapid depletion of the tagged proteins, and their levels were undetectable within an hour of auxin addition (Figure 1C-D, Supplementary Figure S1G-H).

**Figure 1.**
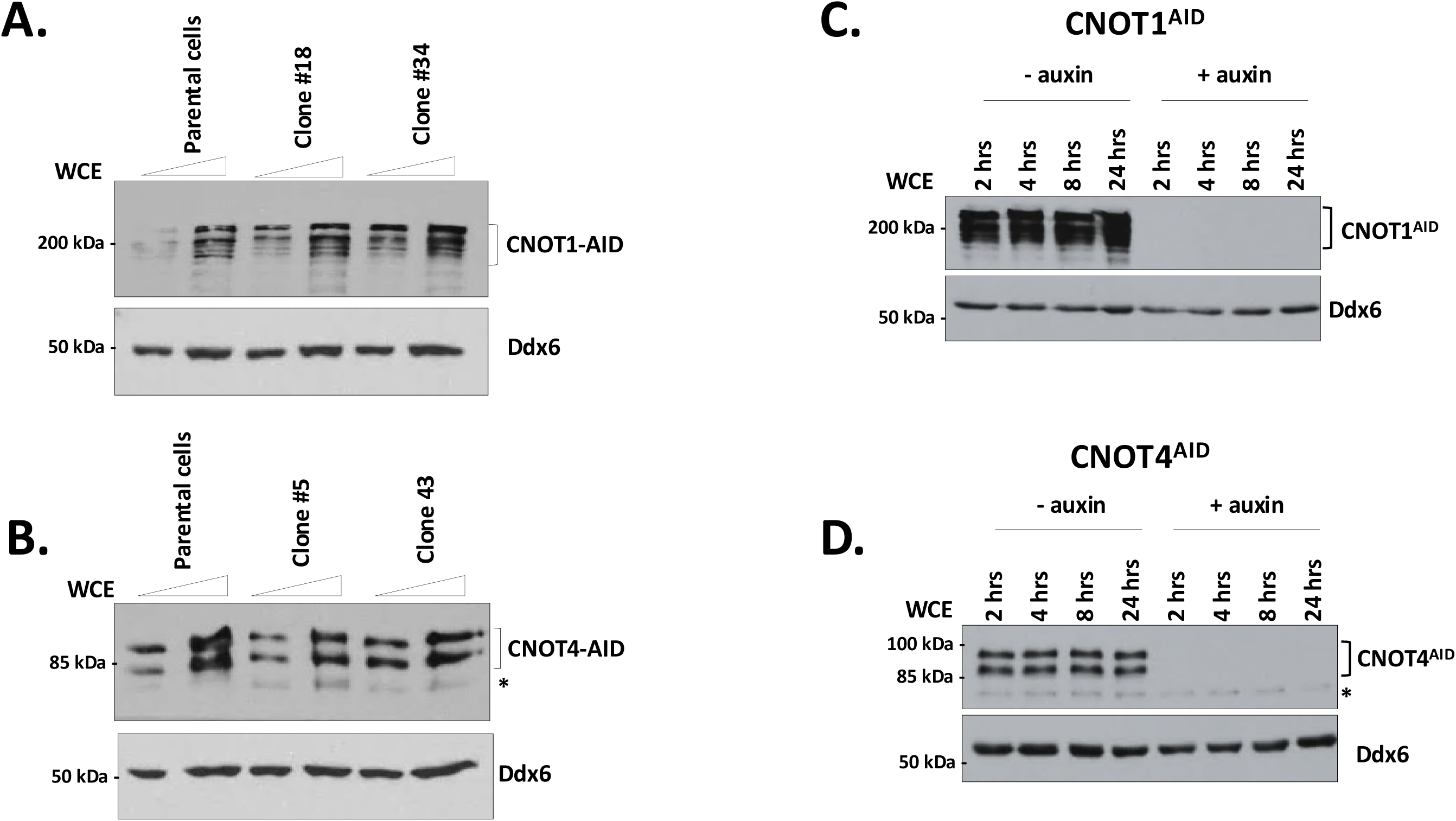
Auxin inducible degron system to deplete CNOT1 and CNOT4. **(A, B)** Western blot analysis of parental (DLD-1^TIR1+^), CNOT1^AID^ and CNOT4^AID^ cells. Two clones were analyzed. Two different amounts of lysates (differing by a factor of 2) were loaded onto SDS-PAGE followed immunoblot analysis using antibodies against CNOT1 in (**A)** or CNOT4 in **B).** Ddx6 was used as a loading control. CNOT1^AID^ or CNOT4^AID^ cells were treated with ± 1mM auxin containing media for 2, 4, 8 and 24 h (C and D). Multiple bands detected in CNOT1 blots may represent degradation products or versions with posttranslational modifications. The bands in CNOT4 blots may represent alternate splice isoforms and/or isoforms with alternate translation start sites. The asterisk (in **B** and **D**) represents a non-specific band detected by anti CNOT4 antibody.

siRNA mediated partial knockdown of CNOT1 reduced cell growth, induced cell cycle arrest, and ultimately apoptotic cell death^36,37^. We examined the effects of rapid depletion of CNOT1 and CNOT4 on cell growth and the cell cycle. Depleting CNOT1 suspended cell growth by 24 h and increased death afterwards (Supplementary Figure S2A). In contrast, depletion of CNOT4 had no detectable effect on cell growth. However, prolonged auxin treatment (72 h) slowed cell growth in the parental cell line and CNOT4-AID cells (Supplementary Figure S2A). Flow cytometry revealed that depleting CNOT1 for 24 h resulted in an increase of cells in G1/S and a corresponding decrease in cells in the S and G2 phases, but no cell cycle arrest was observed in CNOT4-AID cells (Supplementary Figure S2B). Given these data, analysis of depleted cells was conducted within 24 h.

### Analysis of the effects of subunit depletion on the integrity of CCR4-NOT

Deleting the genes of subunits of the yeast Ccr4-Not complex or knocking down subunits of the human complex by siRNA treatments reduced the accumulation of other subunits^36–38^. To check the effects of CNOT1 and CNOT4 depletion on the complex, we measured the levels of its subunits and monitored complex integrity by size exclusion chromatography (SEC). Depleting CNOT1 decreased the deadenylase subunits, CNOT6 and CNOT7, and CNOT9 protein within 4 h, while CNOT2 and CNOT3 levels did not change. Interestingly, the accumulation of CNOT4 protein increased between 8-24 h of CNOT1 depletion (Supplementary Figure S3A), which we attribute to the stabilization and accumulation of CNOT4 mRNA (data not shown). In contrast, depleting CNOT4 did not affect the levels of any of the subunits examined (Supplementary Figure S3B).

Yeast Not4 stably associates with the Ccr4-Not complex, while CNOT4 does not co-purify with the complex^3,9,39^. However, a prior proximity labeling (BioID) study^40^ suggested it may do so. We conducted a BioID study on CNOT1 and CNOT4 to directly address if CNOT4 interacts with the complex in cells, and we indeed found that CNOT4 biotinylates other CCR4-NOT subunits (Supplementary Figure S4, Supplementary Table S1). We assessed the integrity of the complex using size exclusion chromatography (SEC) on lysates. The CCR4-NOT complex from undepleted cells co-eluted in fractions corresponding to a molecular weight of 1 - 1.5 MDa (Figures 2A and B), which is comparable to the size reported by others^9^. The vast majority of CNOT4 migrated at its calculated MW of ∼66Kda. The very weak signal for CNOT4 in the fractions where the complex elutes was not affected by depleting CNOT1 (Figure 2A), suggesting that this fraction of CNOT4 is not associated with CCR4-NOT. Depleting scaffold subunit CNOT1 resulted in the disassembly of the complex (‘+auxin’ panels in Figure 2A), as evidenced by the elution of its subunits in lower molecular weight fractions (Figure 2A and not shown). In contrast, depleting CNOT4 did not affect the integrity of the complex, which still eluted at 1-1.5MDa. These data indicate that depletion of CNOT1, but not CNOT4, leads to disassembly of CCR4-NOT, followed by decreased protein levels of many subunits.

**Figure 2.**
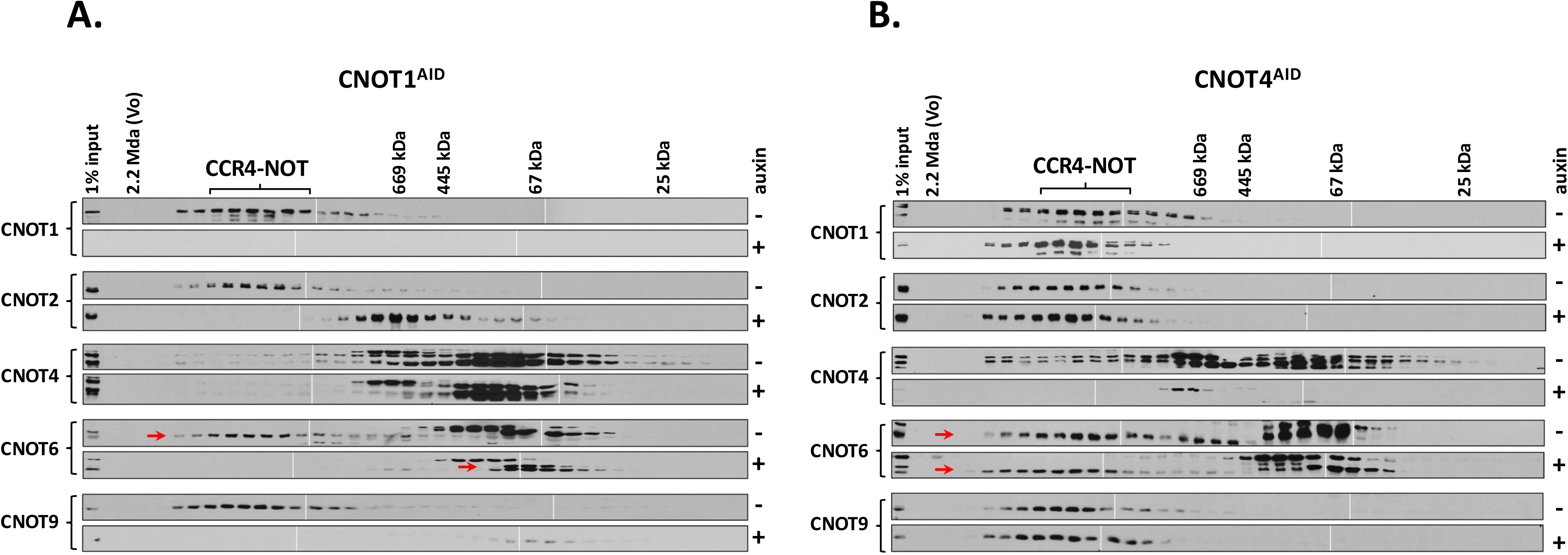
Depletion of CNOT1, but not CNOT4 leads to the destruction of the CCR4-NOT complex. Benzonase nuclease-treated extracts from CNOT1^AID^ (**A**) or CNOT4^AID^ (**B**) cells cultured in ± 1mM were subjected to size exclusion chromatography on a Superose 6 column. The fractions were concentrated by TCA precipitation and subjected to immunoblotting for CCR4-NOT subunits. The red arrow on CNOT6 blot indicates the specific band. Fractions containing the CCR4-NOT complex are indicated above. Molecular weights of calibration proteins are shown on top. For each blot against a particular subunit, multiple blots were spliced together to cover all fractions. White vertical lines represent those splicing junctions. (Vo = the void volume).

### Depletion of CNOT1 and CNOT4 has distinct effects on steady-state RNA levels

Parental cells (DLD-1^TIR1+^), CNOT1^AID^ and CNOT4^AID^ cells were treated with auxin for 2-, 8- and 24-h and the number of genes changing ±auxin at each time point meeting an arbitrary cutoff (FC ≥ 1.5, P^adj^ < 0.01) was calculated. Since CCR4-NOT plays a broad role in mRNA stability, ERCC-spike-in control RNAs were used to normalize the data. The biological replicates of each condition showed a high correlation and introducing the tags into the protein or auxin treatment did not cause gene expression changes in the DLD-1 cells (Supplementary Table S2 and Supplementary Figure S5A-B). Depleting CNOT1 led to predominantly increased RNA abundance of a significant number of genes by 2 h, and the number increased over time (Figure 3A-C, top panels for CNOT1^AID^, Supplementary Figure S5C, Supplementary Table S3). This was expected considering CNOT1’s role in mRNA deadenylation [see below and^1,36,41^]. Greater than 90% of the differentially expressed genes (DEGs) were protein-coding genes (Supplementary Figure S5D). In contrast to what was observed in CNOT1-depleted cells, depletion of CNOT4 showed relatively mild effects on steady-state levels of RNAs (Figure 3A-C, bottom panels for CNOT4^AID^ and Supplementary Figure S5C). After an initial appearance of a handful of DEG at 2 h, the number reduced to essentially zero at 8 h, followed by another very modest increase of ∼650 DEG at 24 h. We suspect this complex pattern is due to compensatory effects on transcription at 8 h and secondary effects at 24 h (see below). The differences in RNA abundance changes in CNOT1-versus CNOT4-depleted cells may be explained in part by the effects of the depletion of each protein on complex integrity. Depleting CNOT4 maintained the nuclease activities of CCR4-NOT while depleting CNOT1 destroyed the complex (Figure 2). Gene ontology (GO) analysis on the DEGs in CNOT1-depleted cells returned terms related to transcriptional regulation (Supplementary Figure S6A). Depletion of CNOT4 for 2 h indeed resulted in DEGs that were involved in transcriptional regulation as well, but later time points resulted in no significant enrichment (24 h depletion) of these terms, and but instead genes involved in cell cycle regulation showed significant enrichment (Supplementary Figure S6B). This was unexpected since CNOT4-depleted cells showed no cell cycle arrest or growth defects (Supplementary Figure S2B). Further, genes involved in cell cycle regulation were indeed enriched in the DEGs of long-term CNOT1 depletion (24 h) (enrichment FDR = 5.7E-31), which was consistent with the cell cycle arrest phenotype in these cells (Supplementary Figure S2B). Although, there was a large difference in the number of DEGs in CNOT1- and CNOT4-depleted cells (Figure 3), there was a substantial and significant overlap in the DEGs at the 2 and 24 h time points (Supplementary Figure S6C-D).

**Figure 3.**
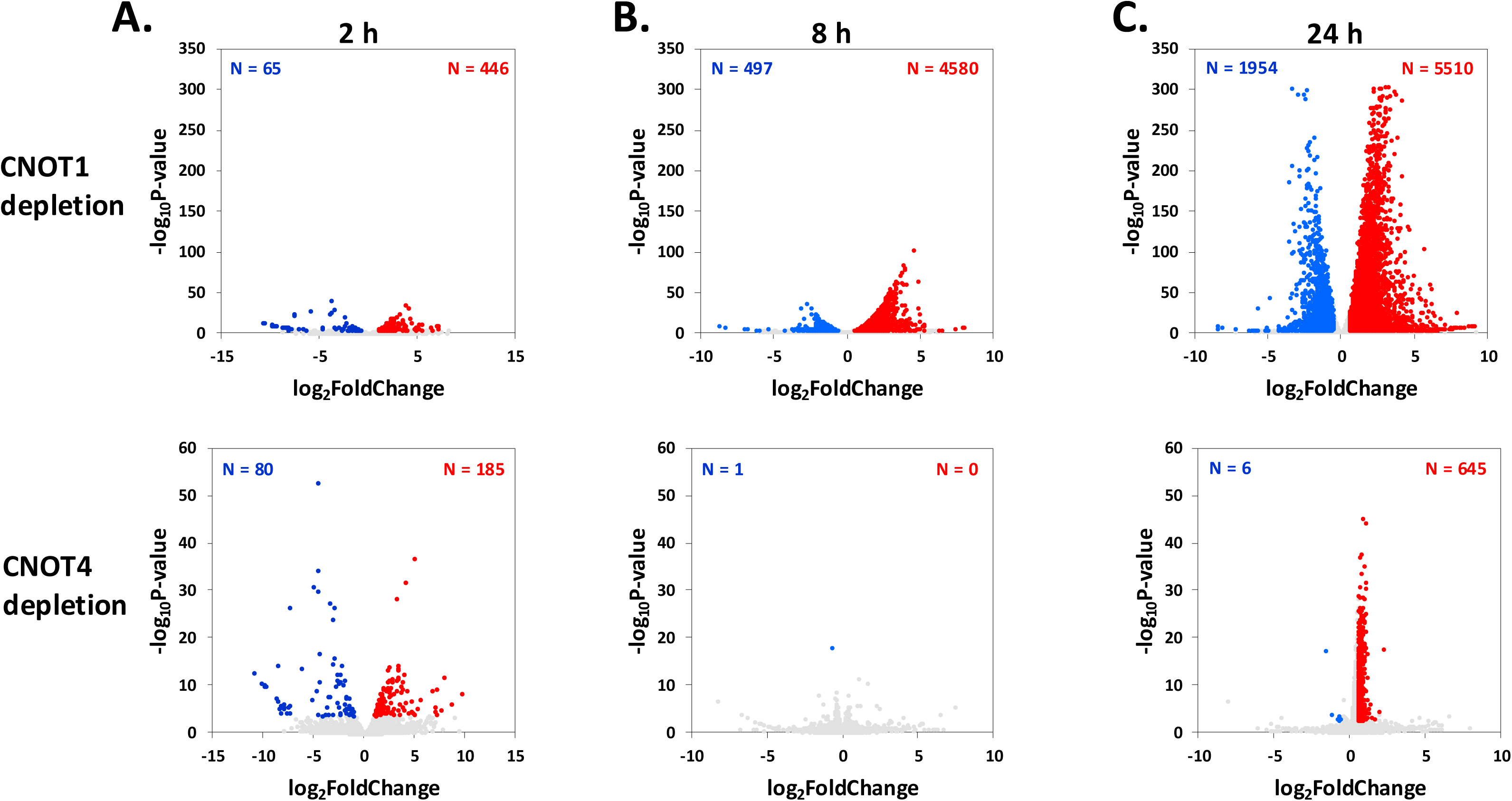
Effects of CNOT1 and CNOT4 depletion of steady-state RNA levels. **(A, B, C)** Volcano-plots of log_2_FoldChange (FC) in steady-state RNA levels versus negative log_10_ of p-values. CNOT1 or CNOT4 were depleted with 1mM auxin for 2 h (**A**), 8 h (**B**) and 24 h (**C**). The numbers in each panel represent differentially expressed genes with a FC ≥ 1.5 and p-adj < 0.01. For 2 h datapoint FCs were calculated by comparing RNA levels between auxin treated CNOT1^AID^ cells or CNOT4^AID^ cells (depleted) versus auxin treated parental cells (non-depleted). For **A, B and C**, each dot represents an individual RNA.

### Unexpected opposite effects of CNOT1- and CNOT4-depletion on RNA stability

We measured the effects of depleting each subunit on mRNA stability. Transcription was inhibited using Triptolide (TPL), which has been shown to inhibit overall transcription by targeting TFIIH^42^ and has been used to calculate RNA half-lives transcriptome-wide^43^. The levels of RNAs were measured at different time points (3, 6 and 12 h) and RNA half-lives were calculated from the data (Figure 4A, Supplementary Table S4). First, we confirmed that the estimated half-lives in parental (un-depleted) cells correlated well with previously published datasets (Supplementary Figure S7A). Consistent with CCR4-NOT’s role in deadenylation, depleting CNOT1 resulted in a global stabilization of RNAs (Figure 4B), where the median half-life increased from 10.6 h to 24.3 h (Figure 4D). Surprisingly, depleting CNOT4 increased the RNA turnover rate (Figure 4C), reducing the median half-life to 6.5 h (Figure 4D). The scatter plot from the CNOT4 depletion displayed a distinct pattern. We speculate that this is explained by the decay capacity that has plateaued-mRNAs can’t be turned over any faster due to a limited factor(s). Cumulative fraction analysis further confirmed these findings (Figure 4E). Although CNOT1 and CNOT4 are subunits of the same complex, their depletion conferred opposite effects on mRNA stability. The effects on RNA half-lives were predominantly seen in protein coding messenger RNAs (Supplementary Figure S7B). To rule out that the changes in half-lives were caused by reduced fitness or metabolic changes in the cells, we examined the changes in half-life of non-polyadenylated histone mRNAs, which should be insensitive to the inactivation of CCR4-NOT^44^. Indeed, depletion of either CNOT1 or CNOT4 had minimal effect on the turnover of non-polyadenylated histone mRNAs (Figures 4F, Supplementary Figure S7C-D). *H1.0* is a polyadenylated histone mRNA, and its stability changed like other mRNAs (Supplementary Figure S7E). These results were validated by RT-qPCR analysis of specific messages (Supplementary Figure S8A-I).

**Figure 4.**
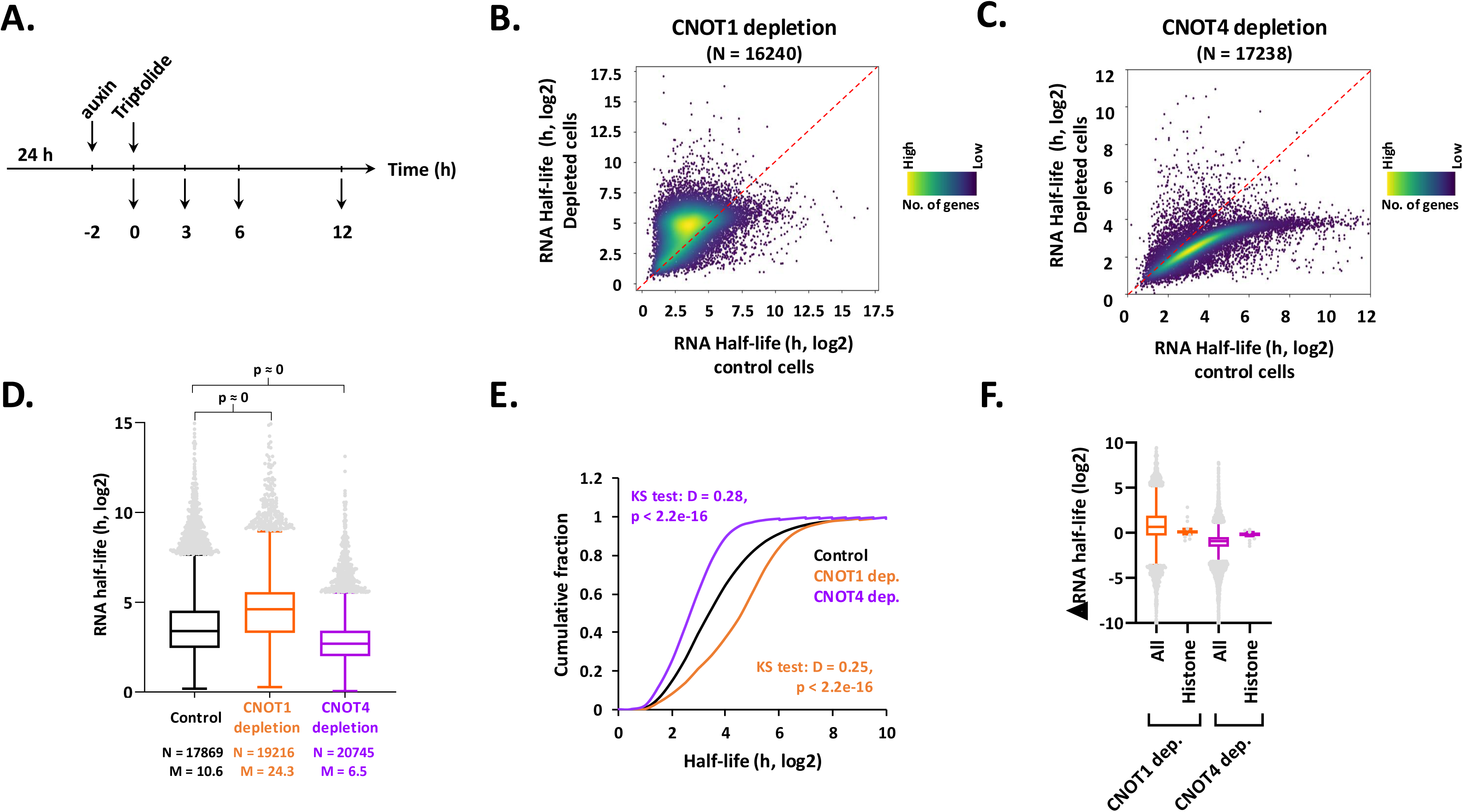
Depletion of CNOT1 and CNOT4 have opposite transcriptome-wide effects on RNA half-lives. **(A)** Schematic depicting the experimental design to determine RNA half-lives. **(B)** Scatter plot of RNA half-lives calculated in control (parental DLD-1^TIR1+^ cells +auxin) and CNOT1 depleted cells (CNOT1^AID^ +auxin). The biological replicates of the RNA-seq data showed high correlation (R > 0.93, Supplementary Table S2). **(C)** Same as panel B except that data from CNOT4 depleted cells (CNOT4^AID^ +auxin) are plotted. **(D)** Boxplot analysis of RNA half-lives. p-values were calculated using the Mann Whitney Test between CNOT1 and CNOT4 depleted cells versus control cells. Total number of datapoints (N) and median half-life (M, in h) are shown **(E)** Cumulative fraction distribution plots of RNA half-lives in control and CNOT1 and CNOT4 depletion conditions. Kolmogorov-Smirnov (KS) statistic is shown. **(F)** Boxplot analysis of change in RNA half-lives upon CNOT1 and CNOT4 depletion for all and non-polyadenylated histone mRNAs. ‘All’ correspond to all RNAs whose foldchange in RNA half-life could be calculated under the experimental conditions (N = 16240, for CNOT1 depletion; N = 17238 for CNOT4 depletion). Sixty-one non-polyadenylated histone mRNAs were included in the analysis and polyadenylated histones were excluded.

### Effects of depletion of CNOT1 and CNOT4 are influenced by codon optimality content and steady-state turnover rates

To address whether the changes in RNA turnover are related to the steady state half-life of the RNA, we sorted RNAs into quartiles based on half-life. Highly unstable RNAs (Q1) showed the greatest increase in half-life (median increase in half-life ∼3-fold) in CNOT1-depleted cells. However, the most stable RNAs (Q4) showed a significant decrease in half-life compared to all RNAs (Supplementary Figure S9A). The destabilizing effect of CNOT4 depletion on RNAs was observed in all the quartiles, although highly stable RNAs in Q4 (t_1/2_ > 22.7 h) showed the strongest decrease in half-lives (median reduction in half-life ∼ 4.5-fold) (Supplementary Figure S9B). This confirms the opposite effects of CNOT1-versus CNOT4 depletion on RNA decay.

Since codon optimality affects mRNA turnover^45–47^ and CCR4-NOT senses codon optimality and targets mRNAs with a high number of non-optimal codons for degradation^8^, we asked if the sensitivity of mRNAs to CCR4-NOT depletion was related to their codon optimality. We used two datasets of codon optimality, codon stability coefficients [‘CSC’,^45^] and partial least squares score for codon optimality [‘PLS’,^48^]. First, we confirmed that the half-lives we calculated under control (undepleted) conditions correlated significantly with these optimality matrices (Spearman’s rho ≥ 0.33, p ≤ 10^-73^) (Supplementary Figure S9C, D), supporting the previously known relationship between codon content and mRNA half-lives. We hypothesized that if CCR4-NOT regulates mRNA turnover depending upon codon optimality, the mRNAs enriched with non-optimal codons would be more sensitive to CNOT1 depletion. Indeed, the non-optimal mRNAs (CSC-lowest or PLS-lowest) showed a greater increase in half-lives in CNOT1-depleted cells than their counterparts (CSC-highest or PLS-highest) (Supplementary Figure S9E, F). These findings underscore the role of CCR4-NOT in regulating mRNA half-lives as a function of codon optimality. The mRNAs with the lowest optimality content showed a relatively lesser decrease in half-lives in CNOT4-depleted cells as compared to mRNAs with the highest optimality content (Supplementary Figure S9G, H), suggesting that non-optimal short-lived mRNAs are less sensitive to CNOT4 depletion than optimal long-lived mRNAs. Thus, although codon optimality correlated with changes in half-life in both CNOT1- and CNOT4 depletion conditions, they responded oppositely.

### Depletion of CNOT1 and CNOT4 leads to wide-spread increase in nascent transcription

Yeast Ccr4-Not regulates transcription by promoting elongation and controlling TATA-binding protein^2,3,49^, but it is unclear if human CCR4-NOT does so. We performed transient transcriptome sequencing (TT-seq) using 4-thio-uracil labeling to measure ongoing transcription^50^. DLD-1^TIR1+^, CNOT1^AID^ and CNOT4^AID^ cells were treated with auxin for 2- and 8-h, followed by a 15-minute pulse of labeling (Supplementary Figure S10A). Data was normalized to a spike-in control (*S. pombe* 4-thioU labeled RNA, Supplementary Table S5). Interestingly, depleting CNOT1 or CNOT4 significantly increased nascent transcription across the genome within 2 h (Figure 5A and Supplementary Figure S10B). Protein depletion takes at least 1 h (Figure 1 and S1), so the increased transcription was very rapid. Thousands of differentially transcribed genes (DTGs) were identified (Figure 5A and Supplementary Figure S10C-F), of which ≥ 85% of the genes showed enhanced transcription (> 1.5 FC, p^adj^ < 0.01 cutoff) and were predominantly protein-coding mRNAs (∼90% of DTGs, (Supplementary Figure S10G-I). The fast response to depletion (2 h) on transcription and the opposite effects of depleting CNOT1 versus CNOT4 on mRNA half-lives, suggests the increased transcription is not an indirect effect of altered mRNA decay. Although, we suspect the stabilization of mRNAs in CNOT1-depleted cells could explain why the number of differentially transcribed genes (DTGs) in CNOT1-depleted cells increased from ∼800 at 2 h to ∼6500 at 8 h of depletion, whereas that in CNOT4-depleted cells did not change much (∼1200 at 2 h and ∼1600 at 8 h). Since the depletion of CNOT1 results in both increased transcription and decreased decay, there was a global increase in steady-state RNA levels, including those coding for transcriptional regulators (Supplementary Figure S6). This might lead to increased levels of transcription regulators, causing a further increase in nascent transcription at 8 h in the CNOT1-depleted cells.

**Figure 5.**
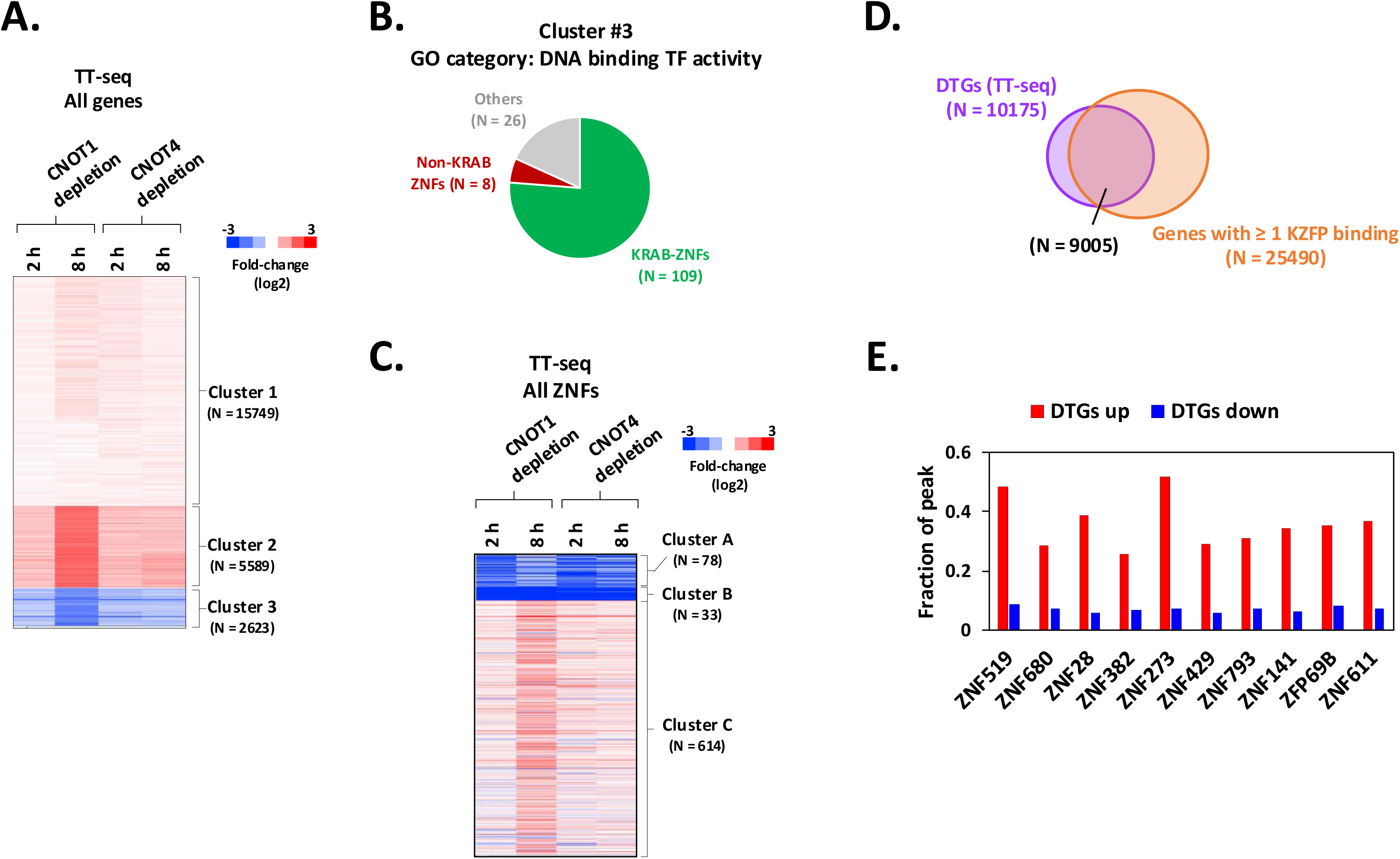
Depletion of CNOT1 or CNOT4 leads to wide-spread increased nascent transcription and decreased expression of KRAB Zinc Finger genes on chromosome 19. **(A)** K-means clustering of fold-changes (FCs) in RNA transcription of all genes calculated by TT-seq. The rows represent individual RNAs (N = 23961). FC data were used to make heatmaps in ‘Cluster 3.0’ (K = 3 and ‘Euclidean distance’ as a similarity metric) and were visualized by ‘Java TreeView’. The biological replicates showed high reproducibility (R> 0.99, Supplementary Table S2). **(B)** Distribution of genes in the top enriched GO category in Figure S11B (N = 143). ZNFs were grouped as with or without KRAB domain, based on^35^. **(C)** K-means clustering of FC in transcription of ZNF genes determined by TT-seq (N= 725). Heatmap was generated as described in **A**. **(D)** Venn diagram showing overlap of DTGs (Figure S10C-F) and genes showing at least one KZFP binding peak. Data from repressed KZFPs with highest number of genome-wide peaks were analyzed. **(E)** Analysis of fraction of KZFP peaks in genes which are transcriptionally either ‘Up’ or ‘Down’ in CNOT1/4 depletion conditions.

### CNOT1 and CNOT4 regulate transcription of the KRAB-Zinc finger repressor family genes

K-means clustering of the transcriptomics data showed a widespread increase in transcription in CNOT1- and CNOT4-depleted cells (Figure 5A, clusters 1 and 2). However, the genes in cluster 3 were notably reduced (Figure 5A, cluster 3). GO analysis of genes in cluster 3 produced terms related to ‘DNA binding transcription factor activity’ (Supplementary Figure S11A. 143 genes, 2.1x enrichment, 2.3E-14 FDR). Most of the transcription factors in this cluster (109/143) were KRAB-ZNFs genes (Kruppel associated box containing Zinc finger protein coding, hereafter referred to as KZNFs), which function in gene repression (Figure 5B, Supplementary Figure S11B-C). To gain more insights into this novel finding, we clustered the transcription changes of all ZNF genes in the dataset (N = 725). Most ZNF genes increased in depleted cells, like the rest of the genome (Figure 5C, cluster C, N = 614), but the transcription of KZNF genes (N = 102) was notably reduced (Figure 5C, clusters A and B, Supplementary Figure S11C).

KZFPs bind throughout the genome, and we speculated that the widespread increase in nascent transcription is correlated with reduced KZFP expression. We used KZFP Chip-exo data^34^ to see if the KZNFs in cluster 3 (Figure 5A) bind near genes with increased transcription. We found that ∼90% of the DTGs had at least 1 KZFP binding peak within or ±1KB of the gene (Figure 5D). The binding peaks of the repressed KZFPs were more enriched at genes whose transcription increased than those that were repressed by CCR4-NOT depletion (Figure 5E). This strong association suggests that reduced KZFP expression contributes to the widespread increase in transcription in CCR4-NOT depleted cells.

### CNOT1 and CNOT4 depletion increases transcription of repeat elements, particularly LINEs

KZFPs genes cluster on chromosome 19, and they expanded and diversified in response to the rise of novel endogenous retroelements^28,34,51,52^. Most KZNFs repressed by CCR4-NOT inactivation are on chr19 (Figure 6A). KZFPs bind to and repress endogenous retroviral elements and protein coding genes by binding to promoters and ancient LTRs adjacent to genes^20,26,34,35,53^. To test if depleting CNOT1 and CNOT4 derepressed repetitive elements, the TT-seq data was remapped using the Allo tool, which efficiently allocates multi-mapped reads to locations within the genome and is capable of mapping reads to repeat elements^54^. The data revealed that the expression of LINE retrotransposons (LINEs) and other repeat elements, including some SINEs and LTRs of endogenous retroviruses was increased (Figure 6B, C, Supplementary Table S6). We then analyzed the expression of the mapped LINEs (N = 15611) and found a striking increase in LINEs transcription within 2 h of depleting CNOT1 or CNOT4 (Figure 6D, clusters 1 and 2, ∼77% of all LINEs analyzed) Furthermore, the timing of LINE activation coincided with KZNF gene repression (Figure 5C, clusters A and B). At 8 h of CNOT1 and CNOT4 depletion, the global de-repression of LINEs expression was still observed (Figure 6E, cluster 2 and 4), but there was a subset of LINEs that were repressed in CNOT1-depleted cells (cluster 1). The recovery of LINE repression at 8 h could be explained by the increased steady-state levels of some KZNF mRNAs due to reduced mRNA decay, therefore re-establishing repression by recovering KZFP levels (Figure 4 and data not shown).

**Figure 6.**
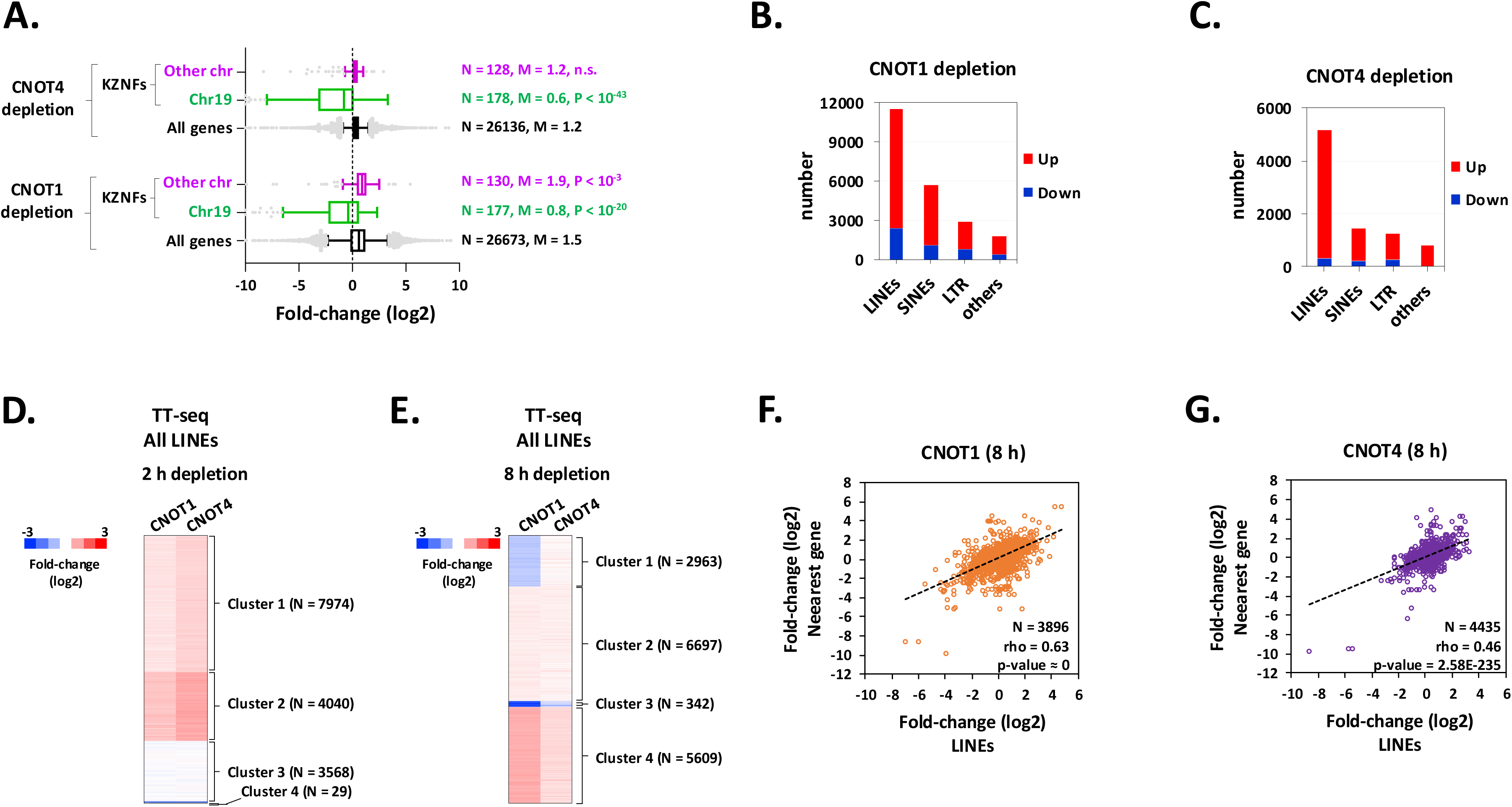
Depletion of CNOT1 or CNOT4 leads to activation of transcription of LINEs and other repeat elements. **(A)** Boxplot analysis of changes in transcription of KRAB-ZNFs (KZNFs) genes after 8 h depletion of CNOT1 or CNOT4. KZNFs were binned based on their location on chromosome 19 or other chromosomes based on^35^. Total number of datapoints (N), median fold-change (M) and P-values by Mann Whitney Test (compared to All) are indicated in the panel. **(B, C)** Distribution of differentially transcribed repeat elements, rTEs, (FC > 1.5 and p^adj^ < 0.05) in CNOT1-(B) and CNOT4-depleted cells (**C**). The fold-change in expression was determined by DESEq2 analysis of RNA expression measured by TT-seq as described in Figure S10B; and the differentially transcribed rTEs at either 2 h or 8 h depletion were plotted. ‘Others’ include DNA transposons and a minor class of repeat elements. **(D, E)** K-means clustering of fold-changes in transcription of LINEs upon depletion of CNOT1 or CNOT4 for 2 h (**D**) or 8 h (**E**). **(F, G)** Correlation plots between fold-change in transcription (TT-seq) of LINE elements versus that of its nearest gene upon 8 h depletion of CNOT1 (**F**) or CNOT4 (**G**). Top ten KZFPs with highest number of genome-wide peaks based on^34^ and whose transcription was repressed upon CNOT1 or 4 depletion were analyzed (as in Figure 5D-E). LINEs associated with these KZFPs were mapped and the nearest gene was located for each LINE followed by correlation analysis. Number of datapoints (N), Spearman correlation (rho) and 2-sided p-value are shown on each plot.

LINEs impact the expression of adjacent genes by controlling chromatin, acting as an enhancer, or providing an alternate promotor and transcriptional start site (TSS)^20,27,34,53^. To investigate if changes in transcription of RNAs are linked to LINE expression, we correlated the change in transcription of KZFP-bound LINEs with that of the adjacent gene. We focused on the KZFPs with the highest number of peaks genome-wide, which were repressed in CCR4-NOT depleted cells (the same set of KZFPs was analyzed in Figure 5D-E). The analysis revealed a highly significant positive correlation between the change in LINE transcription and that of their adjacent genes (Figure 6F-G). Similar correlations were observed with other transposable repeat elements, such as SINEs and LTRs. (Spearman’s rho > 0.37, p-value < 8.42E-85, data not shown). Together, these findings suggest that CCR4-NOT ability to suppress transcription genome-wide is related to its functions in regulating KZFPs and LINEs.

## Discussion

CCR4-NOT has been thoroughly studied in yeast, where it was initially defined as a transcription regulator and later was found to carry out other post-transcription functions [reviewed in^1–3^]. However, much less was known of human CCR4-NOT functions, especially in transcription. Our work fills this gap and makes the remarkable discovery that it silences repeat elements and genome-wide transcription, at least in part, by maintaining the expression of KZNF genes (Figure 7).

**Figure 7.**
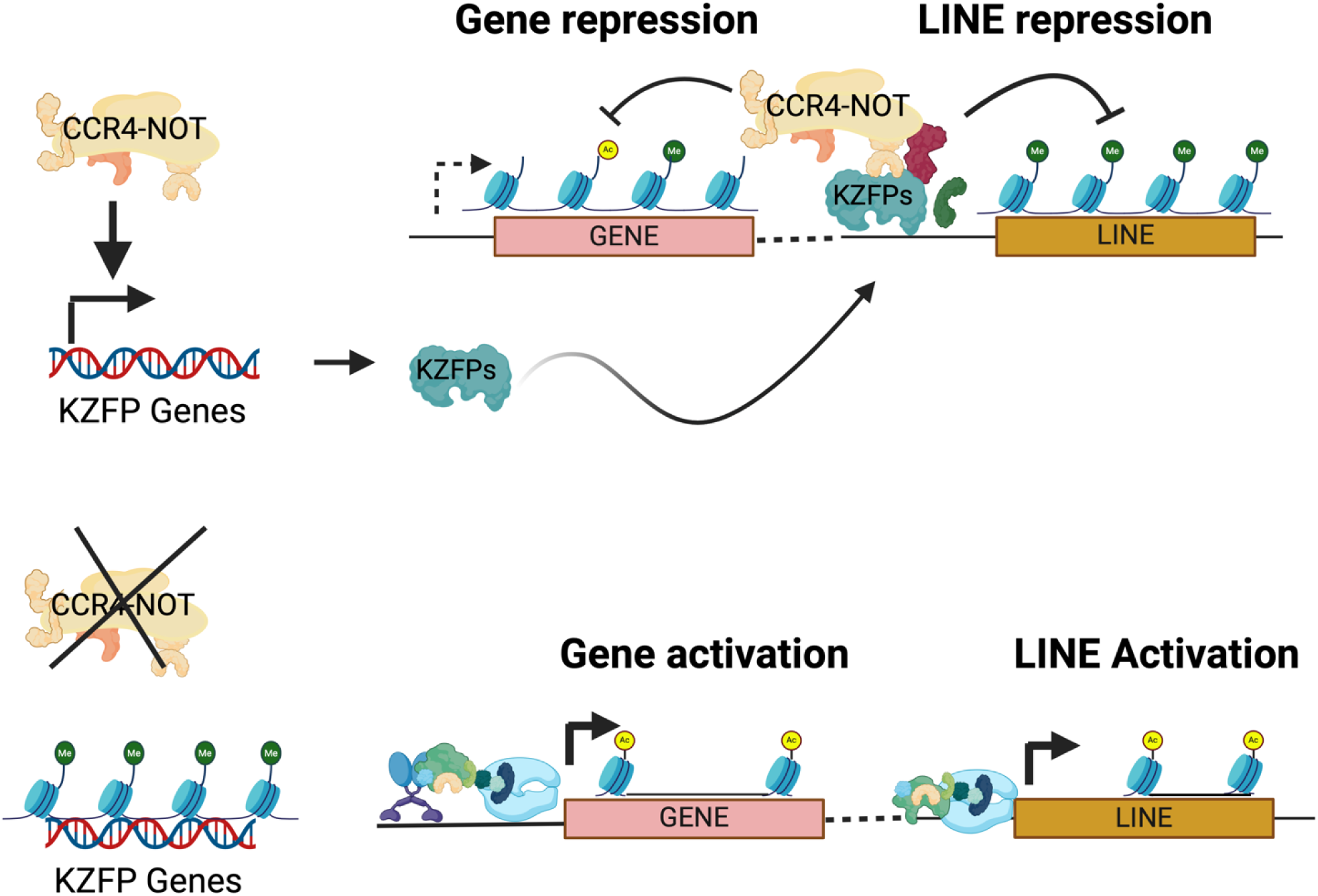
Model for the regulation of gene expression by CCR4-NOT complex. CCR4-NOT maintains normal gene expression by driving KZFP expression and directly repressing LINEs. Targeting both KZNF expression and LINEs directly achieves tighter repression or provides a redundant mechanism to silence rTES (top). Loss of CCR4-NOT reduces KZFP expression, causing depression of repeat elements and adjacent genes (bottom).

We addressed the roles of CNOT4 in mRNA metabolism, which had not been reported. In contrast to what was observed when CNOT1 was depleted, depleting CNOT4 resulted in a quite remarkable increase in mRNA decay rates (Figure 4). RNF219 is another RING domain-containing E3 ligase that associates with CCR4-NOT and, like what we found for CNOT4, knocking it down accelerated mRNA decay^55^. RNF219 binds to the CNOT9 module, like CNOT4. This begs the question of whether different subcomplexes exist, one with CNOT4 and another containing RNF219. The authors of the prior study speculated that RNF219 attenuates deadenylase activity by impeding the access of poly-A RNA to the catalytic module or by reducing the activity of the deadenylases in CCR4-NOT. The function of CNOT4 can be envisaged in a similar context. However, there are two other explanations for this phenotype. First, CNOT4 could regulate the partitioning of mRNAs into distinct pools in the cell. For example, yeast Ccr4-Not regulates the solubility of mRNAs, which impacts their degradation. The solubility of mRNAs correlates with the codon optimality and co-translational decay, which are inversely regulated by Not1 and Not4^56^. Second, the enhanced decay in CNOT4-depleted cells is the response to the increased synthesis of mRNAs through transcript buffering. Transcript buffering is a phenomenon where a change in synthesis is balanced by an opposing compensatory change in decay to maintain steady-state RNA levels^57–59^. As stated above, little change in steady-state RNA levels was observed in CNOT4-depleted cells in spite of increased synthesis (Figures 3 and 5). The same response was not observed in the CNOT1-depleted cells because this condition destroyed the CCR4-NOT complex (Figure 2) and, specifically, the major deadenylase activity in cells (Figure 4 and Supplementary Figure S3). The machinery needed to respond to the increased transcription to enact buffering is inactive; thus, CNOT1-depleted cells lose the capacity to buffer the change in synthesis. In contrast, depleting CNOT4 did not affect the integrity of the CCR4-NOT complex (Figure 2), enabling buffering of the increased nascent transcription (synthesis).

The most striking outcome of our study is that we identified a novel role for CCR4-NOT in suppressing rTE expression and global gene expression. The activation of rTEs and global transcription in CCR4-NOT depleted cells correlated with the repression of KZNF genes (Figures 5 and 6), and we found a substantial overlap between genes whose transcription increased and the presence of one or more KZFP binding events (Figure 5D). KZFPs are the largest family of transcriptional repressors in vertebrates. The human genome has ∼350 KZNFs; and ∼200 are positioned within six clusters on chromosome 19^28^. KZFPs function in transcriptional silencing by repressing transposable elements (TEs), mainly retrotransposons^26^, such as LINEs. LINEs are a class of non-long terminal repeat (LTR) retrotransposons that have enormous potential for regulating chromatin structure and transcription of adjacent genes^19,31–33,53^. rTEs are widely distributed throughout the genome, located near or within genes. We propose that the widespread increase in canonical gene transcription, at least partly, is linked to the loss of rTE silencing due to reduced KZNF expression (Figure 7), which is consistent with the strong correlation between the rTE activation change and that of the adjacent gene (Figure 6F-G). The precise mechanism of how CCR4-NOT regulates KZNF expression has yet to be determined, but one possibility is that CCR4-NOT directly drives KZNF gene transcription.

The regulation of KZNFs by CCR4-NOT likely contributes to establishing and maintaining rTE silencing throughout cell development and growth. However, this cannot fully explain the rapid, within 2 h, activation of rTEs in CCR4-NOT depleted cells. KZNF mRNA and protein levels would need to drop substantially to relieve repression. One possibility is that CCR4-NOT targets regulatory factors that control KZFPs expression, possibly via the stabilization of a labile regulatory factor by CNOT4-dependent ubiquitination. The other is a parallel, direct repressive effect on rTEs. CNOT3 and TRIM28 were identified as factors required for stem cell renewal and preventing differentiation of mouse stem cells and CNOT3 and TRIM28 binding to promoters overlapped considerably^60^. TRIM28/KAP1 is a corepressor utilized by KZNFs to silence rTE expression^26,34^. CCR4-NOT may regulate TRIM28 and repression at rTEs. Since activation of rTEs is so deleterious to cells, it makes sense that multiple, reinforcing mechanisms have evolved to prevent their activation.

Spurious activation of rTE, especially LINEs, causes genome instability and contributes to disease^21,24,25^. CCR4-NOT in yeast and higher eukaryotes is required to maintain genomic integrity^1,37,61,62^. Recently, it was shown that partially knocking downing CNOT1 by siRNA transfection increased bulk transcription, suggested by 5-ethynyl uridine (EU) incorporation and microscopy of stained cells^37^. This low-resolution method might have picked up the increased rTE transcription we demonstrated here. Interestingly, increased R-loop (RNA:DNA hybrids) formation and genome instability were also observed in the knockdown cells. R-loops are enriched in genomic regions with a higher abundance of L1 LINEs^63^. Our discovery that CCR4-NOT suppresses global transcription and rTE activation suggests a novel mechanism for how gene regulatory complexes preserve genome stability and integrity.

## Methods

### Cell lines

DLD-1 cells (ATCC CCL-221) were purchased from ATCC and DLD-1^TIR1+^ cells were a kind gift from Dr. Mary Dasso at National Institutes of Health, USA. Cells were maintained in complete media [Dulbecco’s Modified Eagle Medium (DMEM) with 10% Fetal Bovine Serum, 1X glutamax and 1X Penicillin and Streptomycin] in 5% CO2 atmosphere at 37 °C in a humidified incubator.

### Plasmid construction

All the primers and plasmids used in this study are listed in Supplementary Table S7. The AID knock-in vectors (donor plasmids) were generated using In-Fusion cloning (Takarabio) generated by PCR. To generate the vector backbone, pVA181 plasmid (see Supplementary Table S7) was digested with EcoRV and NotI and used in *in-fusion* assembly. Clones were screened with restriction enzymes and sequencing. The plasmids for Turbo ID and miniTurboID were generated using In-Fusion cloning as described for mini-AID knock-in vectors, except the sequence coding for mini-AID was replaced with either mini-Turbo-ID; *mTID* Turbo-ID^64^. The sequences coding for mTID and TID were synthesized as gene-blocks from IDT. Plasmid sequences available upon request.

### CRISPR-Cas9 gene editing

A plasmid to produce CNOT1 gRNA or CNOT4 gRNA was made from pX330 (gift from Dr. Mary Dasso, National Institutes of Health) by inserting a double-stranded oligonucleotide corresponding to CNOT1 or CNOT4 gRNA into the BbsI site of the vector. CRISPR mediated mini-AID (referred to AID henceforth) was done as described^65^ with some modification. Cells were seeded in 35 mm dishes so as to reach 50% confluency at the time of transfection. Media were changed to Opti-MEM with 10% FCS. 3 µg of plasmid DNA (1.5 µg of gRNA plasmid + 1.5 µg of donor plasmid) was mixed with 9 µl of Viafect in 300 µl of Opti-MEM. The mixture was incubated at room temperature for 15 min. and was added to cells. 72 h post-transfection, cells were trypsinized, transferred to a 100 mm dish and were cultured in selective complete media with 200 µg/ml Hygromycin-B for 2 weeks. The media containing dead cells were replaced with PBS and colonies were transferred to the wells of a 24-well plate containing trypsin solution, incubated at 37 °C for 15-20 minutes to dissociate cells, followed by addition of fresh complete media with 200 µg/ml Hygromycin. Cells were cultured till formation of monolayer. The cells were cultured in Hygromycin media for one additional passage following which they were cultured in complete media without Hygromycin. Generation of TID/mTID knock-in cell lines was carried out as described for AID tagged cell lines except transfections were done using Lipofectamine3000 using manufacturer’s instructions. Potential clones were screened by PCR screening of gDNA and by western blotting for CNOT1 and CNOT4 (Supplementary Table S7). Details on western blotting and PCR are described in Supplementary Methods.

### Proximity labeling (BioID)

Cells (Parental DLD-1, CNOT1^mTID^, CNOT4^TID^) were seeded in 100 mm dishes for 24 h. At ∼70% confluency old media were replaced with pre-warmed media containing 100 mM biotin. After one hour, media were decanted and cells were washed with ice-cold PBS, collected and lysed in RIPA buffer (50 mM Tris pH 8, 150 mM NaCl, 0.1% SDS, 0.5% sodium deoxycholate, 1% Triton X-100, Protease inhibitors and 1 mM PMSF) on ice for 20 minutes. The lysates were sonicated on Bioruptor^TM^ UCD-200 in ice-water slurry (4 pulses of 30 sec on and 30 sec off on ‘high’ setting). The lysates were clarified by centrifugation at 13,000 rpm at 4 °C for 10 minutes. The supernatant was collected and protein estimation was done using BCA protein estimation kit. To enrich biotinylated proteins, Pierce™ Streptavidin Magnetic Beads (pre-equilibrated with RIPA buffer) were incubated with 1 mg of lysates overnight. The beads were captured on magnet and were subsequently washed in the following order (each wash was 1 ml) - twice with RIPA buffer, once with 1 M KCl, once with 0.1 M Na2CO3, once with 2 M urea in 10 mM Tris-HCl (pH 8.0), twice with RIPA buffer and finally once with 10 mM Tris-HCL, pH 8, 1 mM EDTA. The beads with enriched biotinylated proteins were frozen at −80 °C till further processing. For on-beads tryptic digestion, the beads were washed once with 0.4 ml 50 mM Tris pH 8.5, 2M urea, 1 mM DTT, followed by resuspension in 75 µl 50 mM Tris (pH 8.5), 1 M Urea, 1 mM DTT, and 1 µg of Promega Trypsin Gold Trypsin (0.5 mg/ml prepared in 50 mM acetic acid). The reaction was incubated at 4 h at 37 °C with shaking. The supernatants were collected in fresh tubes and the beads were washed twice with 50 µl 50 mM Tris (pH 8.5), 1 M urea, 1 mM DTT and the washes were pooled with the original elution. DTT was added to a final concentration of 5 mM followed by incubation at 37 °C for 1 h Iodoacetamide was added to this to a final concentration of 10 mM and the reaction was incubated in the dark at room temperature for 30 minutes. An additional 0.5 µg trypsin was added and samples were digested overnight at 37 °C with shaking. The peptides were purified on C18 tips (Pierce-Thermofisher) after acidification with trifluoroacetic acid (TFA) to 0.5% using the manufacturers recommendations. The eluted peptides were dried down and frozen at −20 °C until TMT labelling. The peptides were resuspended in 50 µl 0.1M TEAB buffer followed by addition of TMTpro 16plex reagents and incubated at room temperature for 60 min. The reaction was quenched with hydroxylamine, dried down and repurified on reverse phase tips prior to loading onto Nano-flow Liquid Chromatography (Thermo Easy-nLC 1200). The peptides were loaded on an Acclaim PepMap100 trapping column (75 mm × 2 cm, C18, 5 mm, 100 Å, Thermo), and separated on an Acclaim PepMap RSLC column (50 mm × 15 cm, C18, 2 mm, 100 Å, Thermo) at a flow rate of 300 nL/min in the following gradient of mobile phase B (80% ACN in 0.1% aqueous FA): 5% - 50% B in 210 min, 50% - 90% B in 30 min. A Thermo Orbitrap Eclipse mass spectrometer was operated in a data dependent mode using a method based on the TMT MS2 template. The resulting mass spectra were processed using Proteome Discoverer 3.1 software.

### Chromatographic separation of CCR4-NOT

CNOT1^AID^ or CNOT4^AID^ cells seeded in 100 mm dishes were treated ± 1 mM auxin for 4 h. Cells were collected by scrapping off in ice-cold PBS and lysed in hypotonic lysis buffer (10 mM HEPES, 1.5 mM MgCl2, 10 mM NaCl, 0.2 mM PMSF, 0.5 mM DTT. 1 μg/ml leupeptin, 1 μg/ml aprotinin and 1 μg/ml pepstatin A. 1mM benzamidine-HCL) on ice for 10 min, homogenized and nuclei were pelleted by spinning at 4000 rpm [JS-4.2 rotor], 4 °C, 15 minutes. The supernatant was added with 0.11 volume of 10X cytoplasmic extract buffer (0.3 M HEPES, 1.4 M NaCl, 0.03 M MgCl_2_, protease inhibitors. 0.2 mM PMSF, 0.5 mM DTT) and Benzonase (1 μl per 600 μl extract), incubated on ice 30 minutes. The lysate was further spun down at 13,000 rpm for 20 min, at 4 °C. The supernatant was dialyzed at 4 °C in dialysis buffer (20 mM HEPES/NaOH, 150 mM NaCl, 10 μM ZnCl_2_, 1mM DTT, 10% glycerol, 0.5 mM PMSF, 0.2 μg/ml leupeptin, 0.2 μg/ml aprotinin and 0.5 μg/ml pepstatin A, 1 mM benzamidine-HCL) for 4 hours, spun at 14,000 rpm for 10 min at 4 °C and the supernatant was concentrated using Vivaspin concentrator (20 kDa cutoff) to 5-6 mg/ml protein. Two to three mg of lysate was loaded onto a Superose 6 size exclusion chromatography column, run on ӒKTA pure™ chromatography system (GE Life sciences) in chromatography buffer (20 mM HEPES/NaOH, 150 mM NaCl). Fractions were concentrated by TCA precipitation and analyzed by western blotting.

### RNA extraction

Following the treatment with ±1mM auxin for desired time, media were decanted and the cells were lysed directly in 1 ml Trizol. The cells were resuspended thoroughly by pipetting up and down several times and were incubated at room temperature for 5 minutes. 0.2 ml chloroform was added per 1 ml of Trizol. The tubes were capped and shaken vigorously by hand for 15 seconds and then were incubated at room temperature for 2-3 minutes. The mixture was then centrifuged at > 12,000 xg for 15 min at 4 °C, the aqueous phase was collected in a separate tube followed by addition of equal volume of 100% ethanol. The RNA was cleaned up using Ambion RNA PureLink Kit according to manufacturer’s instructions and eluted in 100 µl RNase-free water.

### DNase treatment, tape station, cDNA synthesis and RT-qPCR

10 µg of total RNA was treated with Turbo DNase (TURBO DNA-free™ Kit) for 30 minutes at 37 °C. 10 µl of DNase inactivation agent was added to the reaction followed by incubation at room temperature for 5 minutes. The tubes were flicked 2–3 times during the incubation period to keep the DNase Inactivation Reagent suspended. The reaction was centrifuged at 10,000 × g for 90 seconds and supernatant was transferred in a new tube. The integrity of RNA was assessed by Agilent TapeStation-4150 at the Genomics Core Facility, Penn State University, USA. RNA samples with RIN score > 9.5 were used for further work. 1 µg of DNase digested RNA was used to make cDNA using RevertAid First Strand cDNA Synthesis Kit according to manufacturer’s instructions using Oligo (dT)18 primer and Random Hexamer primer (mixed 1:1). The cDNA product was directly used in PCR applications or stored at - 20 °C until used. RT-qPCR was performed using PerfeCTa SYBR® Green SuperMix (Quantabio). Two dilutions of cDNA were used in RT-qPCR to ensure the linear range of amplification. For each condition at least 2 experimental replicates were used. The RT-qPCR was run on Agilent Real-time PCR machine (G8830A) and the data were analyzed using Agilent AriaMx and Microsoft excel.

### RNA sequencing

Sequencing libraries for steady-state RNA-seq or triptolide RNA-seq to measure half-lives were generated using Illumina Stranded Total RNA Prep with Ribo-Zero Plus according to manufacturer’s instructions. Briefly, 500 ng of Turbo DNase digested RNA was mixed with 2 µl of 1:200 diluted ERCC Spike-in mix (final volume 11 µl) which was subjected to the first step of library prep (i.e. probe hybridization). The sequencing libraries were analyzed on TapeStation-4150 at the Genomics Core Facility, Penn State University, USA and were quantified using NEBNext® Library Quant Kit for Illumina®. The libraries with multiple indices were pooled and treated with Illumina® Free Adapter Blocking Reagent according to manufacturer’s instructions. Sequencing was carried out at Genomics Research Incubator, Penn State University, USA using NextSeq™1000/2000 P2 Reagents(100Cycles) on NextSeq 2000. FASTQ files were further processed as per the Bulk RNA-seq Data Standards and Processing Pipeline (https://www.encodeproject.org/), using STAR [version - STAR_2.5.1b_modified] and RSEM (RNA-Seq by Expectation-Maximization) tool. The reads were mapped to hg38 (GRCh38) using Gencode-v29 comprehensive gene set, predicted tRNA genes, and the ERCC spike-in controls (ThermoFisher Scientific). Read-counts for each gene generated by RSEM were used for differential expression analysis using DESeq2. Genes with a sum of ≥ 8 across both replicates and conditions were used in DESeq2. The expression data were normalized to ERCC spike in controls using estimateSizeFactors function in DESeq2. Genes showing ≥ 1.5-foldchange at p^adj^ < 0.01 were denoted as differentially expressed.

### Transient transcriptome sequencing (TT-seq)

TT-seq libraries were generated as described previously^50^. Briefly, cells (parental, CNOT1^AID^ and CNOT4^AID^) were grown in 150 mm dishes to reach 70% confluency at the time of auxin treatment (2 h or 8 h). Cells were then labelled by addition of 4-Thiouridine (4TU, final concentration 500 µM) to the media for 15 minutes. All the work involving 4TU was performed under conditions of minimal direct light exposure. Media were discarded and cells were immediately lysed in Trizol. RNA extraction was carried out as described earlier. 200 µg of total RNA was mixed with 5 µg of spike in control (total RNA from *S. pombe* labelled with 4TU, described in Supplementary Methods) in a 250 µl reaction. RNA fragmentation was carried out by addition of 50 µl NaOH (1M) to this reaction (40 minutes on ice). RNA fragmentation was stopped by addition of 200 µl Tris (1M, ph 6.8). The RNA was Phenol/Chloroform extracted and used for *in vitro* biotinylation using MTS-Biotin. RNA was heated at 65 °C for 10 minutes, quick-chilled on ice followed by addition of 1X biotin buffer and MTS-biotin (Biotium). Biotinylation was carried out in dark at room temperature for 40 minutes on a rotator and was followed by Phenol/Chloroform extraction to remove excess biotin. After heating at 65 °C and quick-chilled on ice, RNA was captured on Streptavidin beads (Dynabeads™ MyOne™ Streptavidin C1) at room temperature for 20 minutes. Beads were washed in wash buffer (100 mM Tris-HCl, pH 7.4, 1 M NaCl, 10 mM EDTA, and 0.1% Tween 20; prewarmed at 55 °C) 6 times followed by elution by incubating in 100 mM DTT (twice, 5 minutes each). The pooled eluates were precipitated with in presence of 0.5M NaCl and 0.2 mg/ml glycogen. RNA was estimated by Qubit RNA estimation kit.

Sequencing libraries were generated using Illumina Stranded Total RNA Prep with Ribo-Zero Plus according to manufacturer’s instructions. 300 ng of RNA was heated at 65 °C and quick-chilled on ice followed by addition of EPH3 buffer (from library preparation kit) to get the buffer concentrations right for cDNA synthesis followed directly by first strand synthesis. Human (GRCh38) and *S. pombe* (ASM294v2) genomes were concatenated and reads were mapped to the combined genome using STAR aligner (version 2.7.11a). The raw counts of all reads mapping to the annotated region for each gene (i.e. intron + exon) were obtained using featureCounts. The gene read counts were transformed using variance stabilizing transformation (VST) in DESeq2 R-package. Both normalization and differential expression analysis were performed with the DESeq2 package. We used the *S. pombe* spike in read counts for normalization. Genes showing ≥ 1.5-foldchange at p-adj < 0.01 were denoted as differentially expressed.

### KZFP Analysis

Raw sequencing files were downloaded from NCBI for 58 KZFPs (Supplementary Table S8) identified in cluster 3 (Figure 5A) of our TT-seq analysis and whose binding data were previously available^34^. Reads were trimmed using fastp v0.23.4 on default settings. Bowtie2 v2.5.1 was then used to align the single-end reads to hg38 using the parameter “-k 25” to retain multi-mapped reads. Allo v1.1.1^54^ was used to allocate multi-mapped reads on default settings. Peaks were called using Macs2 on default settings. Gene locations were extracted from the GENCODE v29 gtf file for hg38. A given location for a gene was considered the entirety of the gene body, including introns. The resulting bed file was intersected with the KZFP peak calls (+/- 1000bp) using BEDTools v2.31.1.

### Analysis of repeat elements – TT-seq data

Reads were trimmed using fastp v0.23.4 and aligned using STAR v2.2.1 with the arguments “--outSAMmultNmax 25 --outFilterType BySJout” to retain multi-mapped reads. Allo v1.1.1 was used to allocate multi-mapped reads using the arguments “--read-count --splice”. Following this, featureCounts was used to quantify the transcript counts using the arguments “-- countReadPairs -t exon -g gene_id -M” which retains multi-mapped reads. The annotation file used in featureCounts was constructed using the GENCODE v29 gtf file as well as the annotated repeats from RepeatMasker v4.1.3 for hg38. After read summarization, DEseq2 was used to analyze differentially expressed transposable elements. In this analysis, the scaling factors calculated from the spike-in described above were used for normalization. Additionally, canonical genes were removed from the table before DE analysis. We retained them during the featureCounts step in order to avoid transposable elements that overlapped gene exons.

### Reagents

All the relevant reagents used in this study are listed in Supplementary Table S7.

### Webtools and softwares

All the webtools and softwares used in this study are listed in Supplementary Table S7.

## Data availability

All primary data obtained from RNA-Seq, RNA-seq (Triptolide) and TT-seq have been deposited at the Gene Expression Omnibus (Accession ID: GSE277237). Mass spectrometry data will be uploaded to the MassIVE database.

## Author Contributions

S.K. and C.S. performed the molecular and cell biology experiments. S.K., A.M., A.S., B.G., O.T.A., I.A. and S.M. performed gene mapping and bioinformatics analysis. C.A.K advised on library preparation and sequencing. A.A. provided advice on performing gene editing and AID depletion. S.K. and J.C.R. conceived the experiments and wrote the paper.

## Acknowledgements

We thank members of Reese lab and that of the Penn State Center for Eukaryotic Gene Regulation for their helpful comments and suggestions. We would also like to thank Dr. Tatiana Laremore (Penn State Huck Proteomics and Mass Spectrometry Core Facility, RRID:SCR_024462) for mass spectrometric analyses; and several other core facilities at the Penn State University including Genomics (RRID:SCR_023645), Flow Cytometry (RRID:SCR_024460) and Genomics Research Incubator (RRID:SCR_024530). Mary Dasso (NIH) is recognized for providing DLD-1 Tir1 expressing cells and plasmids. Olivia Geesaman assisted in screening plasmid constructs for the BioID experiments. We also acknowledge Sidharth Ghule and Siqing Wang (CHOP Research Institute) for their advice on data analysis and TT-seq library preparation. This research was supported by funds from National Institutes of Health R35 GM136353 (J.C.R) and R35 GM144135 (S.M.).

## References

1. Chalabi Hagkarim, N. & Grand, R. J. The Regulatory Properties of the Ccr4-Not Complex. Cells 9, (2020).

2. Miller, J. E. & Reese, J. C. Ccr4-Not complex: the control freak of eukaryotic cells. Crit. Rev. Biochem. Mol. Biol. 47, 315–33 (2012).

3. Collart, M. A. The Ccr4-Not complex is a key regulator of eukaryotic gene expression. Wiley Interdiscip. Rev. RNA 7, 438–54 (2016).

4. Pavanello, L., Hall, M. & Winkler, G. S. Regulation of eukaryotic mRNA deadenylation and degradation by the Ccr4-Not complex. Front. cell Dev. Biol. 11, 1153624 (2023).

5. Chen, C.-Y. A. & Shyu, A.-B. Mechanisms of deadenylation-dependent decay. Wiley Interdiscip. Rev. RNA 2, 167–83 (2011).

6. Shirai, Y.-T., Suzuki, T., Morita, M., Takahashi, A. & Yamamoto, T. Multifunctional roles of the mammalian CCR4-NOT complex in physiological phenomena. Front. Genet. 5, 286 (2014).

7. Inada, T. & Beckmann, R. Mechanisms of Translation-coupled Quality Control. J. Mol. Biol. 436, 168496 (2024).

8. Buschauer, R. et al. The Ccr4-Not complex monitors the translating ribosome for codon optimality. Science 368, (2020).

9. Lau, N.-C. et al. Human Ccr4-Not complexes contain variable deadenylase subunits. Biochem. J. 422, 443– 53 (2009).

10. Basquin, J. et al. Architecture of the nuclease module of the yeast Ccr4-not complex: the Not1-Caf1-Ccr4 interaction. Mol. Cell 48, 207–18 (2012).

11. Petit, A.-P. et al. The structural basis for the interaction between the CAF1 nuclease and the NOT1 scaffold of the human CCR4-NOT deadenylase complex. Nucleic Acids Res. 40, 11058–72 (2012).

12. Zhang, Q., Pavanello, L., Potapov, A., Bartlam, M. & Winkler, G. S. Structure of the human Ccr4-Not nuclease module using X-ray crystallography and electron paramagnetic resonance spectroscopy distance measurements. Protein Sci. 31, 758–764 (2022).

13. Keskeny, C. et al. A conserved CAF40-binding motif in metazoan NOT4 mediates association with the CCR4-NOT complex. Genes Dev. 33, 236–252 (2019).

14. Chen, Y. et al. A DDX6-CNOT1 complex and W-binding pockets in CNOT9 reveal direct links between miRNA target recognition and silencing. Mol. Cell 54, 737–50 (2014).

15. Albert, T. K. et al. Identification of a ubiquitin-protein ligase subunit within the CCR4-NOT transcription repressor complex. EMBO J. 21, 355–64 (2002).

16. Aslam, A., Mittal, S., Koch, F., Andrau, J.-C. & Winkler, G. S. The Ccr4-NOT deadenylase subunits CNOT7 and CNOT8 have overlapping roles and modulate cell proliferation. Mol. Biol. Cell 20, 3840–50 (2009).

17. Maryati, M., Airhihen, B. & Winkler, G. S. The enzyme activities of Caf1 and Ccr4 are both required for deadenylation by the human Ccr4-Not nuclease module. Biochem. J. 469, 169–76 (2015).

18. Mostafa, D. et al. Essential functions of the CNOT7/8 catalytic subunits of the CCR4-NOT complex in mRNA regulation and cell viability. RNA Biol. 17, 403–416 (2020).

19. Elbarbary, R. A., Lucas, B. A. & Maquat, L. E. Retrotransposons as regulators of gene expression. Science 351, aac7247 (2016).

20. Thompson, P. J., Macfarlan, T. S. & Lorincz, M. C. Long Terminal Repeats: From Parasitic Elements to Building Blocks of the Transcriptional Regulatory Repertoire. Mol. Cell 62, 766–76 (2016).

21. Mendez-Dorantes, C. & Burns, K. H. LINE-1 retrotransposition and its deregulation in cancers: implications for therapeutic opportunities. Genes Dev. 37, 948–967 (2023).

22. Lander, E. S. et al. Initial sequencing and analysis of the human genome. Nature 409, 860–921 (2001).

23. Graham, T. & Boissinot, S. The genomic distribution of L1 elements: the role of insertion bias and natural selection. J. Biomed. Biotechnol. 2006, 75327 (2006).

24. Beck, C. R., Garcia-Perez, J. L., Badge, R. M. & Moran, J. V. LINE-1 elements in structural variation and disease. Annu. Rev. Genomics Hum. Genet. 12, 187–215 (2011).

25. Dumitrache, L. C. & McKinnon, P. J. Out of LINE: Transposons, genome integrity, and neurodegeneration. Neuron 110, 3217–3219 (2022).

26. Ecco, G., Imbeault, M. & Trono, D. KRAB zinc finger proteins. Development 144, 2719–2729 (2017).

27. Rosspopoff, O. & Trono, D. Take a walk on the KRAB side. Trends Genet. 39, 844–857 (2023).

28. Lukic, S., Nicolas, J.-C. & Levine, A. J. The diversity of zinc-finger genes on human chromosome 19 provides an evolutionary mechanism for defense against inherited endogenous retroviruses. Cell Death Differ. 21, 381–7 (2014).

29. Turelli, P. et al. Interplay of TRIM28 and DNA methylation in controlling human endogenous retroelements. Genome Res. 24, 1260–70 (2014).

30. Fasching, L. et al. TRIM28 represses transcription of endogenous retroviruses in neural progenitor cells. Cell Rep. 10, 20–8 (2015).

31. Liu, N. et al. Selective silencing of euchromatic L1s revealed by genome-wide screens for L1 regulators. Nature 553, 228–232 (2018).

32. Lykoskoufis, N. M. R., Planet, E., Ongen, H., Trono, D. & Dermitzakis, E. T. Transposable elements mediate genetic effects altering the expression of nearby genes in colorectal cancer. Nat. Commun. 15, 749 (2024).

33. Ecco, G. et al. Transposable Elements and Their KRAB-ZFP Controllers Regulate Gene Expression in Adult Tissues. Dev. Cell 36, 611–23 (2016).

34. Imbeault, M., Helleboid, P.-Y. & Trono, D. KRAB zinc-finger proteins contribute to the evolution of gene regulatory networks. Nature 543, 550–554 (2017).

35. de Tribolet-Hardy, J. et al. Genetic features and genomic targets of human KRAB-zinc finger proteins. Genome Res. 33, 1409–1423 (2023).

36. Ito, K., Takahashi, A., Morita, M., Suzuki, T. & Yamamoto, T. The role of the CNOT1 subunit of the CCR4-NOT complex in mRNA deadenylation and cell viability. Protein Cell 2, 755–63 (2011).

37. Hagkarim, N. C. et al. Disruption of the Mammalian Ccr4-Not Complex Contributes to Transcription-Mediated Genome Instability. Cells 12, (2023).

38. Maillet, L., Tu, C., Hong, Y. K., Shuster, E. O. & Collart, M. A. The essential function of Not1 lies within the Ccr4-Not complex. J. Mol. Biol. 303, 131–43 (2000).

39. Albert, T. K. et al. Isolation and characterization of human orthologs of yeast CCR4-NOT complex subunits. Nucleic Acids Res. 28, 809–17 (2000).

40. Youn, J.-Y. et al. High-Density Proximity Mapping Reveals the Subcellular Organization of mRNA-Associated Granules and Bodies. Mol. Cell 69, 517–532.e11 (2018).

41. Pavanello, L., Hall, B., Airhihen, B. & Winkler, G. S. The central region of CNOT1 and CNOT9 stimulates deadenylation by the Ccr4-Not nuclease module. Biochem. J. 475, 3437–3450 (2018).

42. Titov, D. V et al. XPB, a subunit of TFIIH, is a target of the natural product triptolide. Nat. Chem. Biol. 7, 182–8 (2011).

43. Gillen, S. L. et al. Differential regulation of mRNA fate by the human Ccr4-Not complex is driven by coding sequence composition and mRNA localization. Genome Biol. 22, 284 (2021).

44. Marzluff, W. F., Wagner, E. J. & Duronio, R. J. Metabolism and regulation of canonical histone mRNAs: life without a poly(A) tail. Nat. Rev. Genet. 9, 843–54 (2008).

45. Forrest, M. E. et al. Codon and amino acid content are associated with mRNA stability in mammalian cells. PLoS One 15, e0228730 (2020).

46. Bae, H. & Coller, J. Codon optimality-mediated mRNA degradation: Linking translational elongation to mRNA stability. Mol. Cell 82, 1467–1476 (2022).

47. Wu, Q. et al. Translation affects mRNA stability in a codon-dependent manner in human cells. Elife 8, (2019).

48. Medina-Muñoz, S. G. et al. Crosstalk between codon optimality and cis-regulatory elements dictates mRNA stability. Genome Biol. 22, 14 (2021).

49. Reese, J. C. The control of elongation by the yeast Ccr4-not complex. Biochim. Biophys. Acta 1829, 127– 33 (2013).

50. Gregersen, L. H., Mitter, R. & Svejstrup, J. Q. Using TTchem-seq for profiling nascent transcription and measuring transcript elongation. Nat. Protoc. 15, 604–627 (2020).

51. Lupo, A. et al. KRAB-Zinc Finger Proteins: A Repressor Family Displaying Multiple Biological Functions. Curr. Genomics 14, 268–78 (2013).

52. Jacobs, F. M. J. et al. An evolutionary arms race between KRAB zinc-finger genes ZNF91/93 and SVA/L1 retrotransposons. Nature 516, 242–5 (2014).

53. Chuong, E. B., Elde, N. C. & Feschotte, C. Regulatory activities of transposable elements: from conflicts to benefits. Nat. Rev. Genet. 18, 71–86 (2017).

54. Morrissey, A., Shi, J., James, D. Q. & Mahony, S. Accurate allocation of multimapped reads enables regulatory element analysis at repeats. Genome Res. 34, 937–951 (2024).

55. Poetz, F. et al. RNF219 attenuates global mRNA decay through inhibition of CCR4-NOT complex-mediated deadenylation. Nat. Commun. 12, 7175 (2021).

56. Allen, G. et al. Not1 and Not4 inversely determine mRNA solubility that sets the dynamics of co-translational events. Genome Biol. 24, 30 (2023).

57. Timmers, H. T. M. & Tora, L. Transcript Buffering: A Balancing Act between mRNA Synthesis and mRNA Degradation. Mol. Cell 72, 10–17 (2018).

58. Sun, M. et al. Global analysis of eukaryotic mRNA degradation reveals Xrn1-dependent buffering of transcript levels. Mol. Cell 52, 52–62 (2013).

59. Haimovich, G. et al. Gene expression is circular: factors for mRNA degradation also foster mRNA synthesis. Cell 153, 1000–11 (2013).

60. Hu, G. et al. A genome-wide RNAi screen identifies a new transcriptional module required for self-renewal. Genes Dev. 23, 837–48 (2009).

61. Jiang, H., Wolgast, M., Beebe, L. M. & Reese, J. C. Ccr4-Not maintains genomic integrity by controlling the ubiquitylation and degradation of arrested RNAPII. Genes Dev. 33, 705–717 (2019).

62. Gaillard, H. et al. Genome-wide analysis of factors affecting transcription elongation and DNA repair: a new role for PAF and Ccr4-not in transcription-coupled repair. PLoS Genet. 5, e1000364 (2009).

63. Zeng, C., Onoguchi, M. & Hamada, M. Association analysis of repetitive elements and R-loop formation across species. Mob. DNA 12, 3 (2021).

64. Branon, T. C. et al. Efficient proximity labeling in living cells and organisms with TurboID. Nat. Biotechnol. 36, 880–887 (2018).

65. Aksenova, V., Arnaoutov, A. & Dasso, M. Analysis of Nucleoporin Function Using Inducible Degron Techniques. Methods Mol. Biol. 2502, 129–150 (2022).

