## supplemental figures and legends for "Human CCR4-NOT globally regulates gene expression and is a novel silencer of retrotransposon activation"

### Supplemental Figure S1

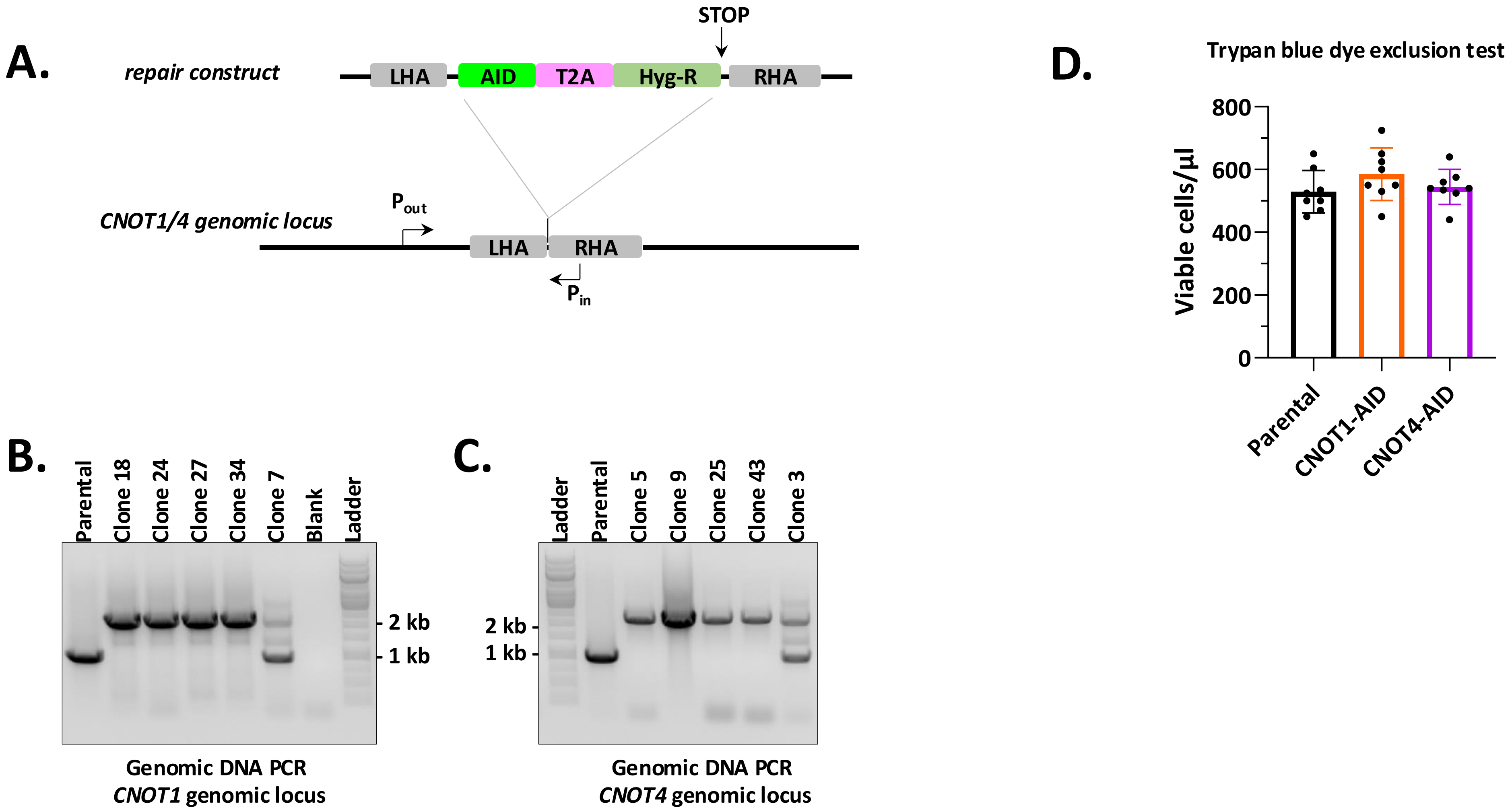

### Supplemental Figure S1

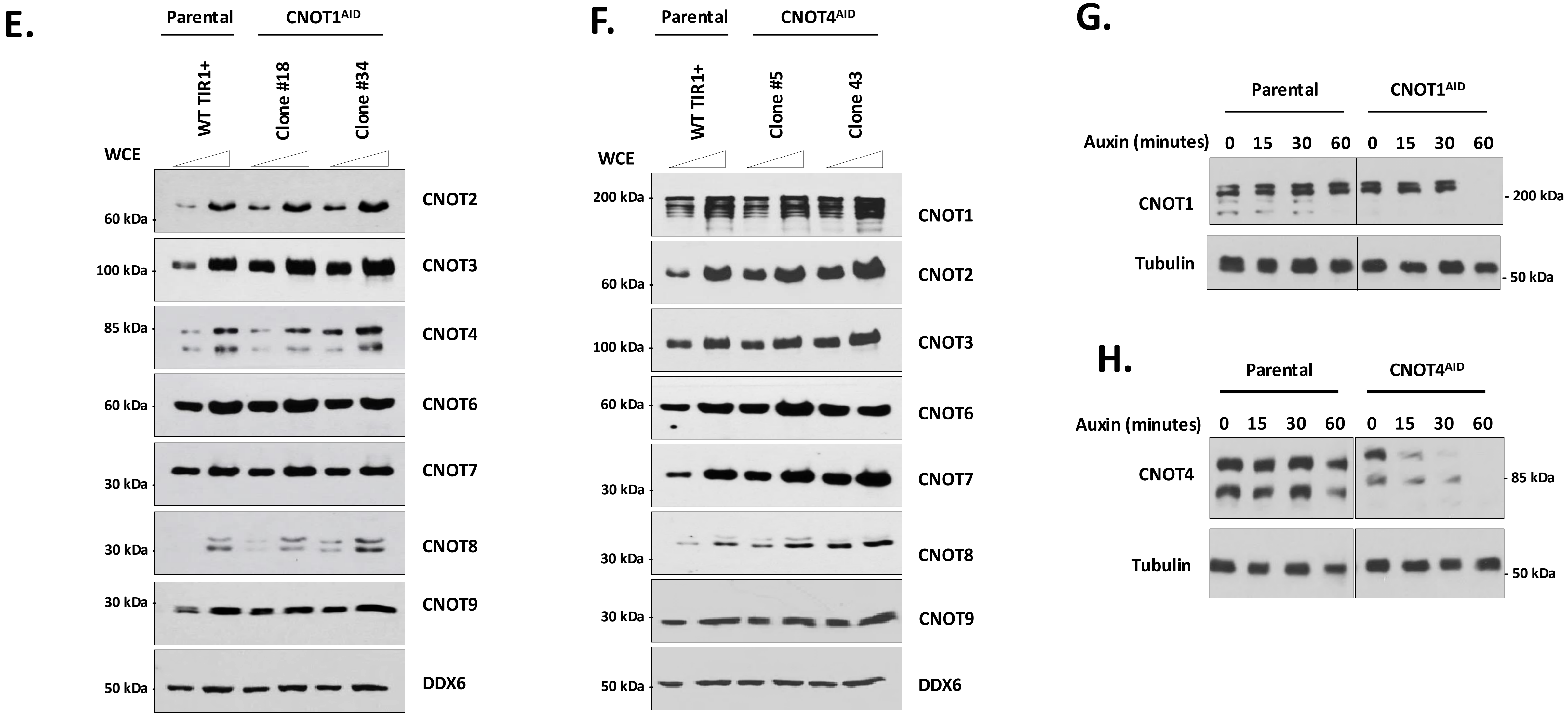

### Supplemental Figure S2

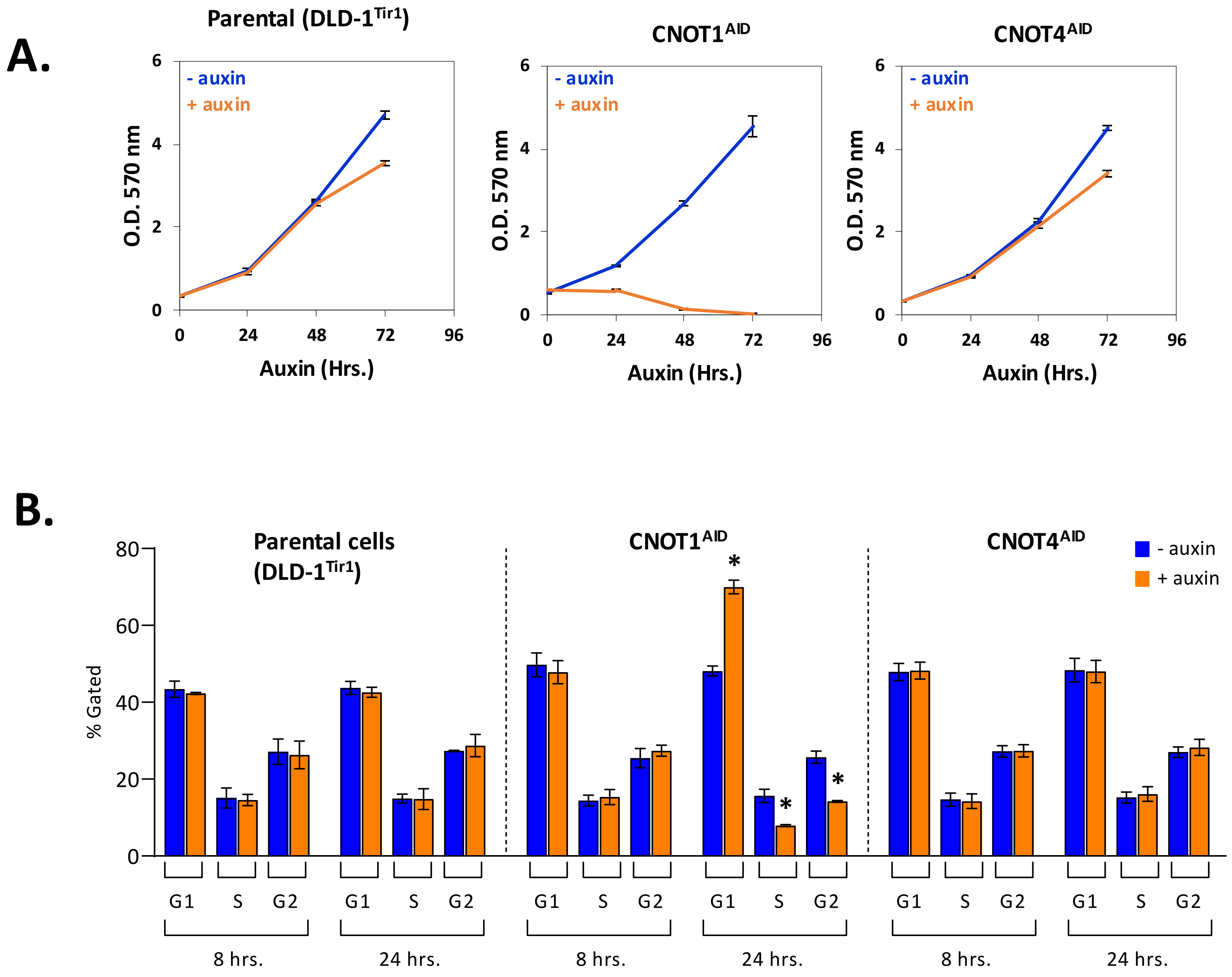

### Supplemental Figure S3

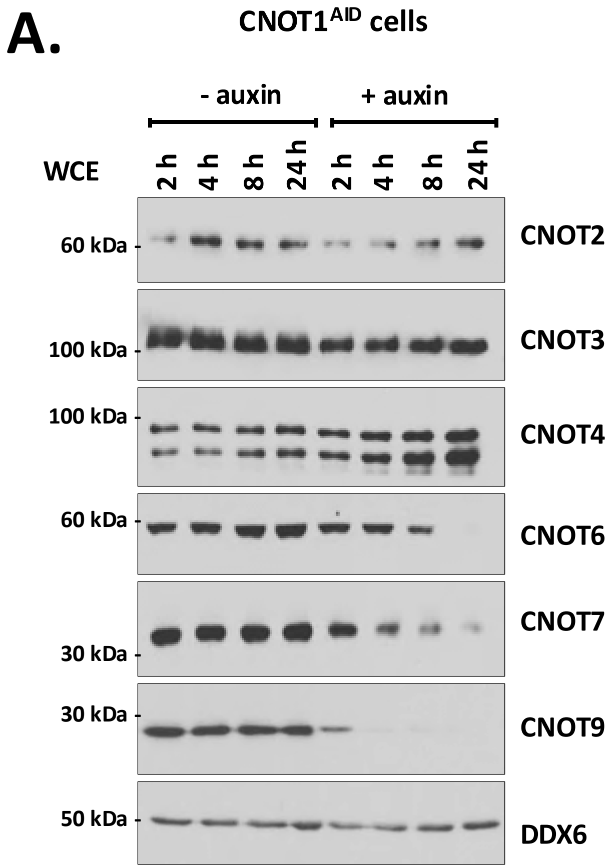


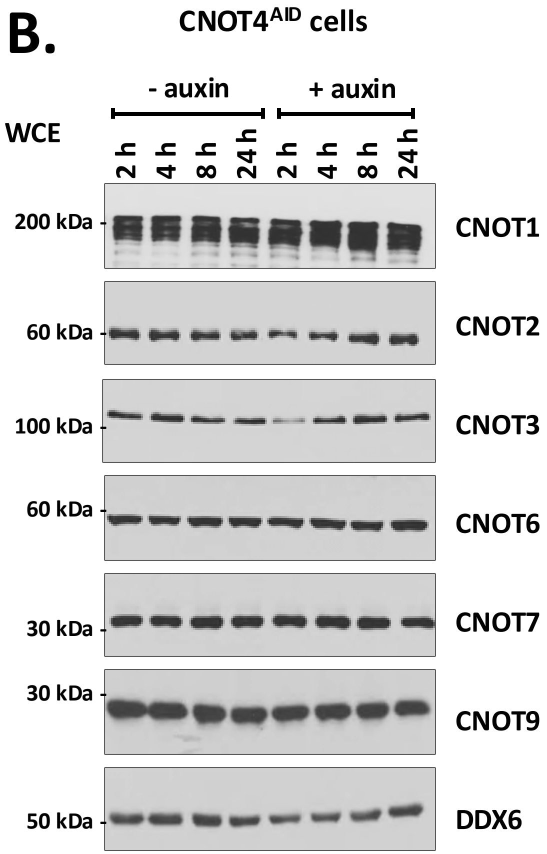

### Supplemental Figure S4

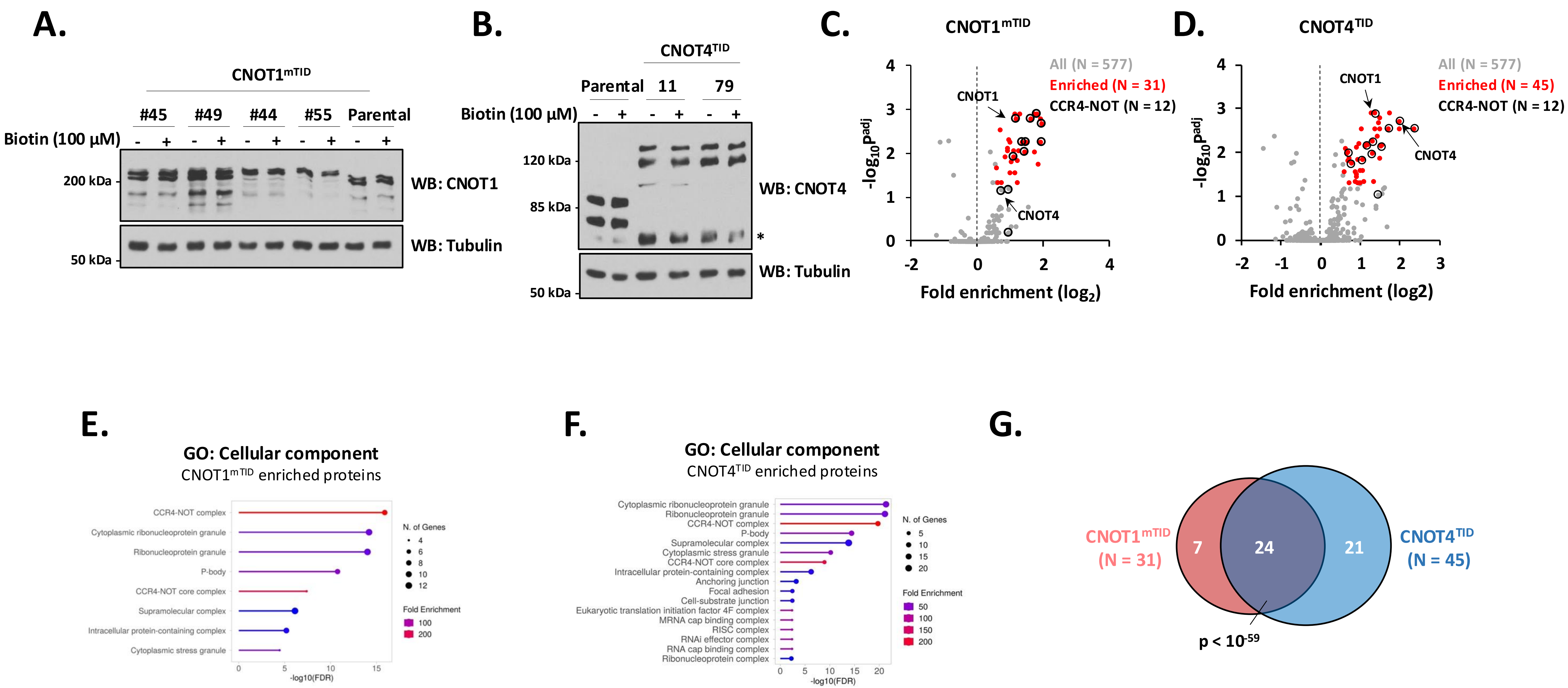

### Supplemental Figure S5

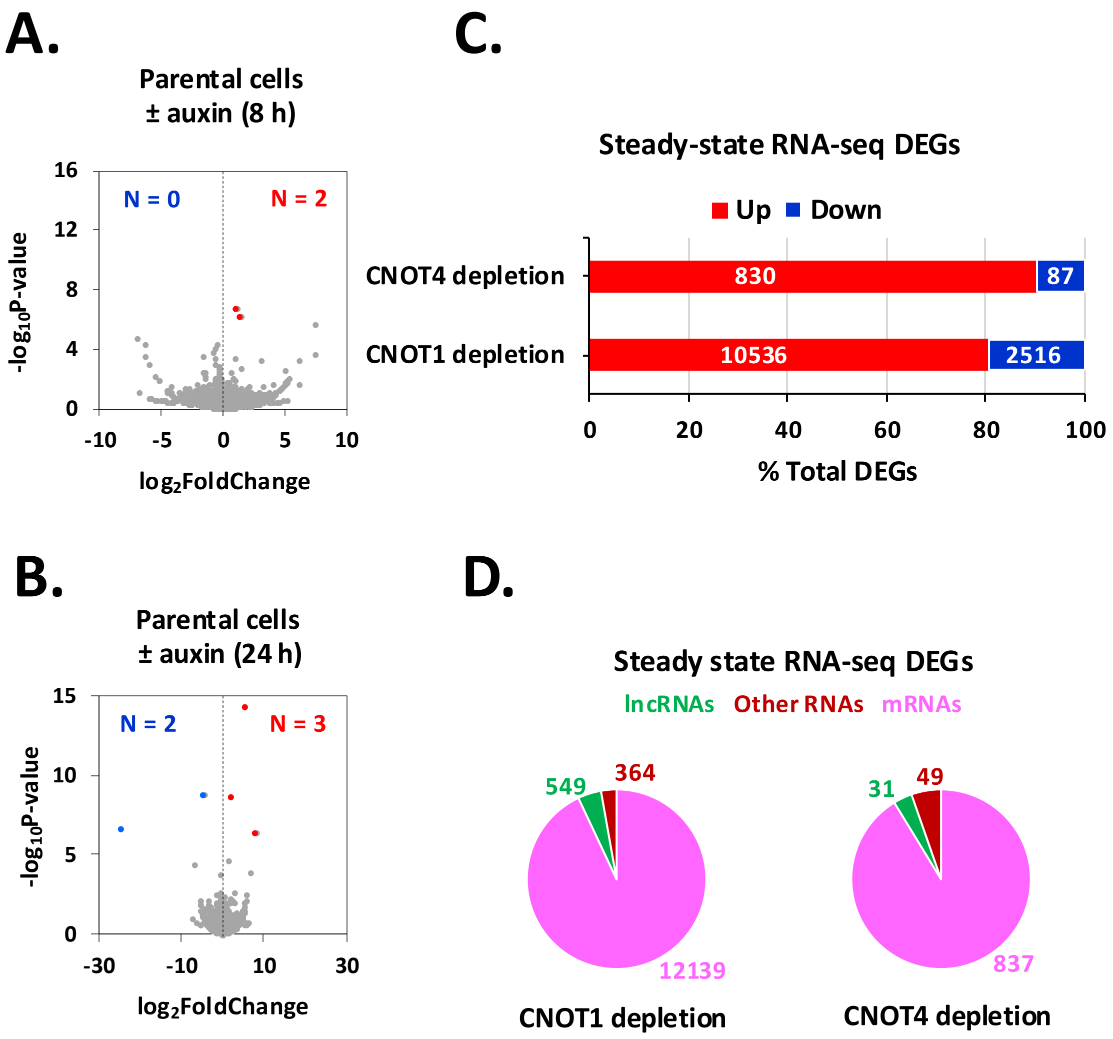

### Supplemental Figure S6

**A.**

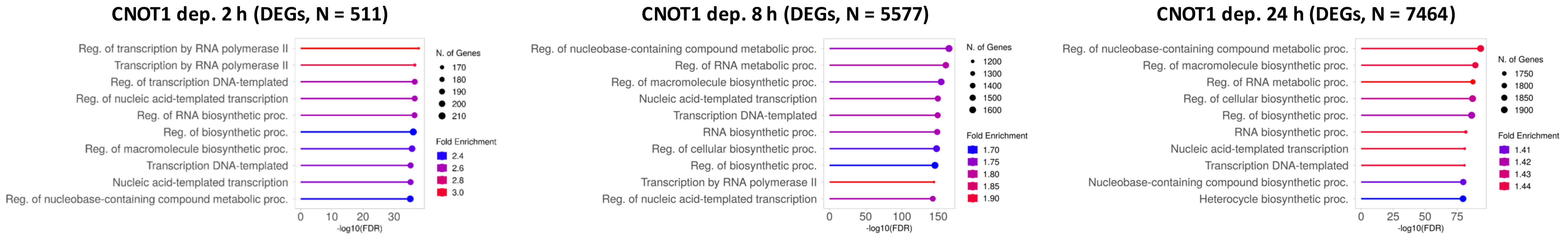


**B.**

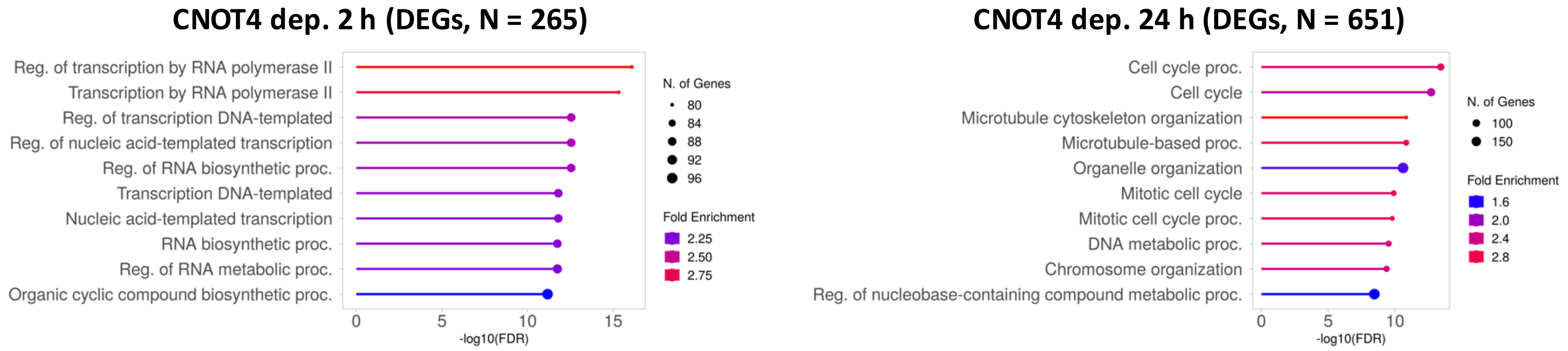


**C.**

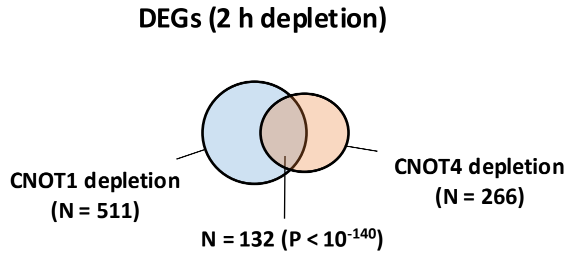


**D.**

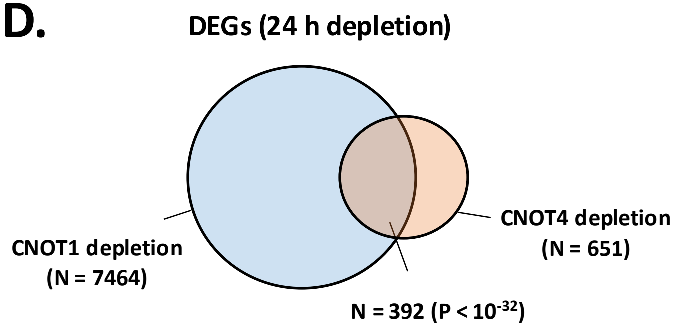

### Supplemental Figure S7

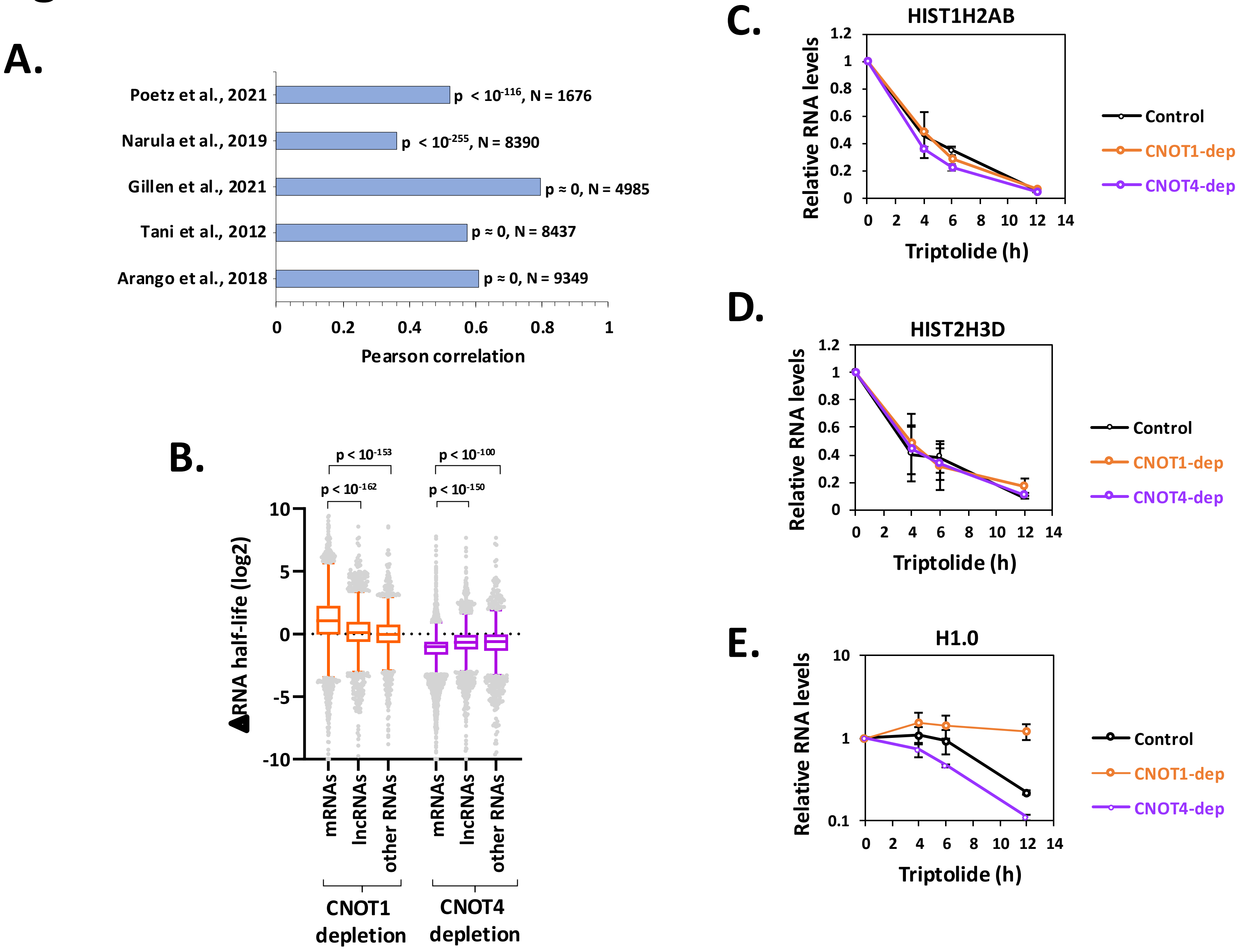

### Supplemental Figure S8

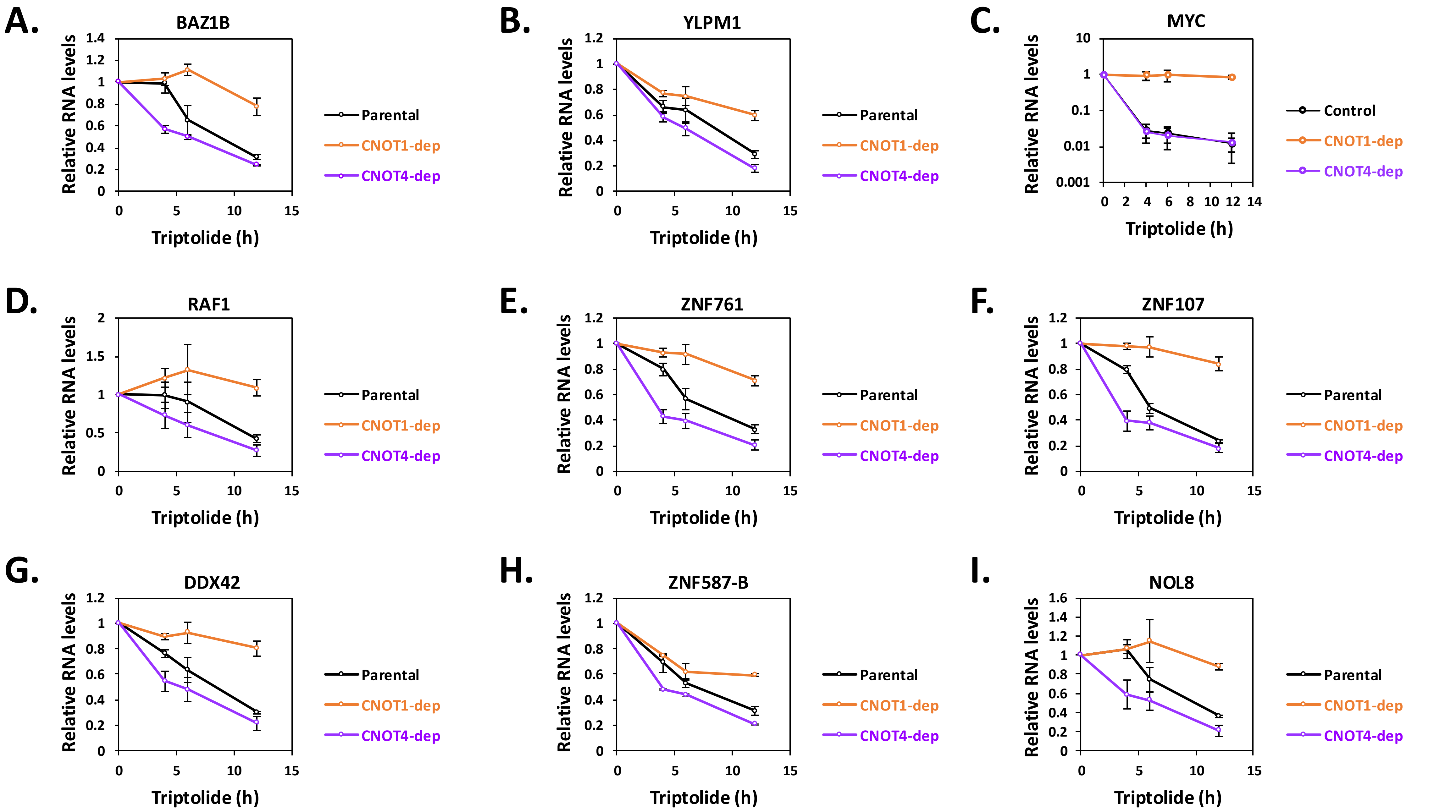

### Supplemental Figure S9

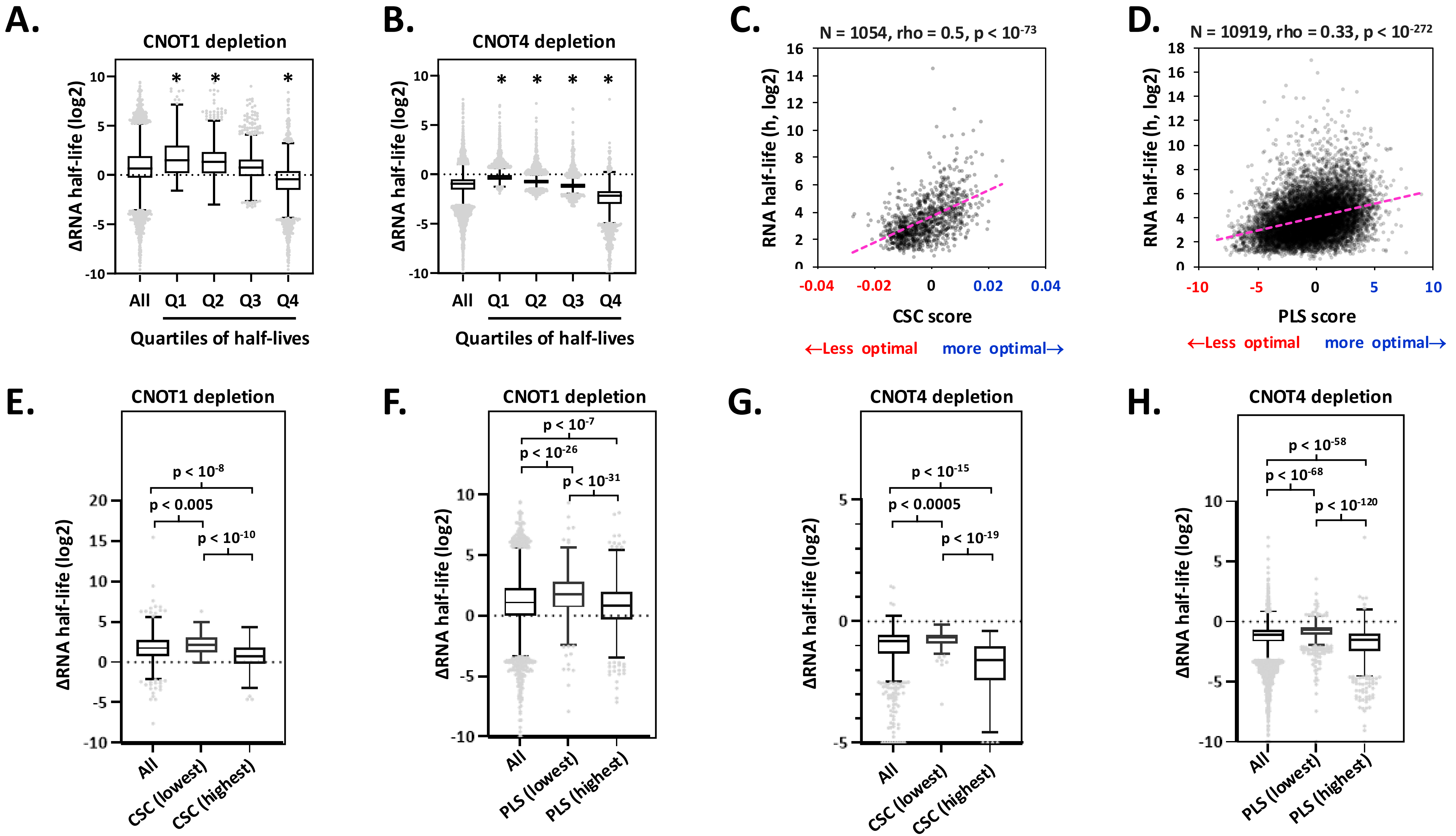

### Supplemental Figure S10

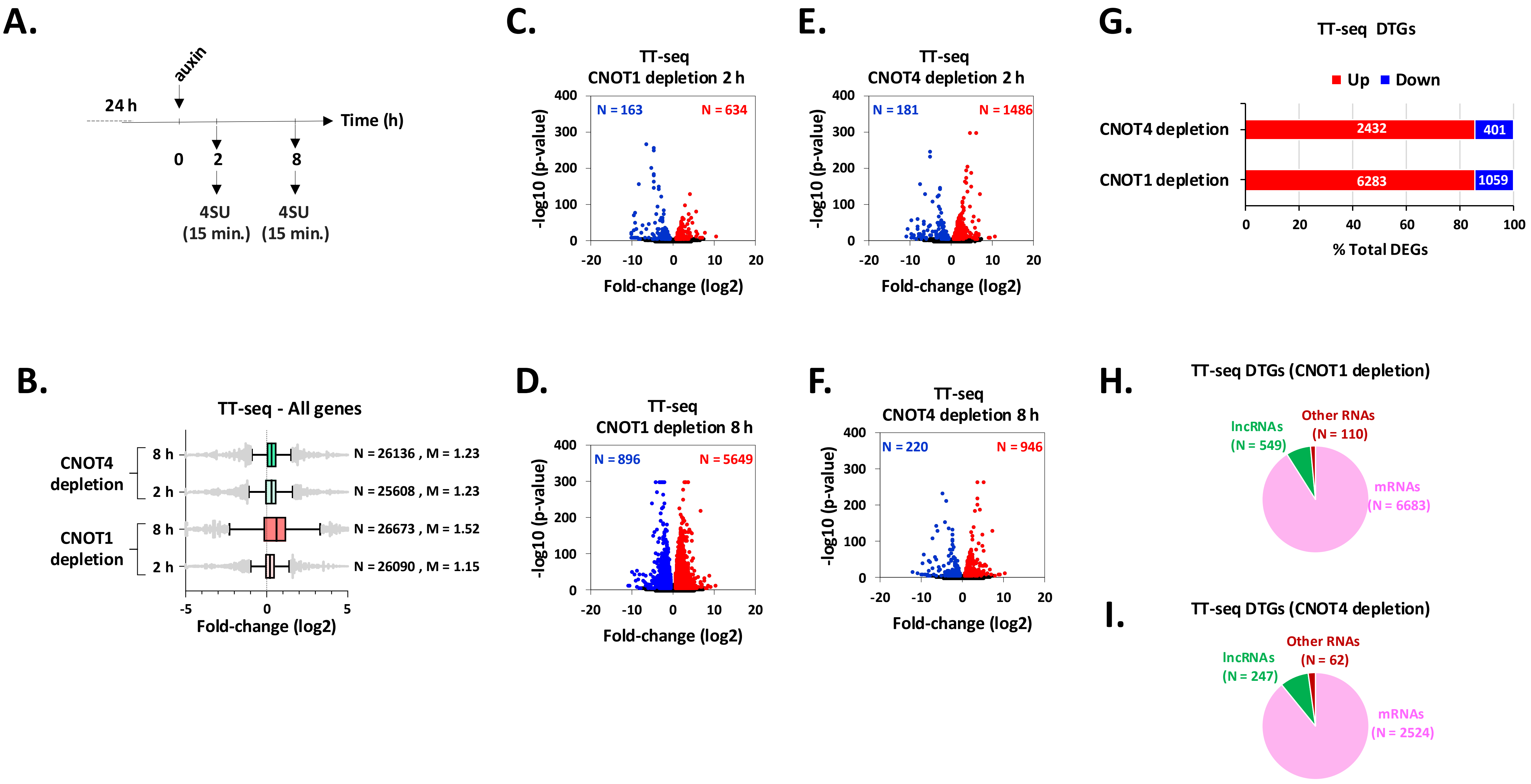

### Supplemental Figure S11

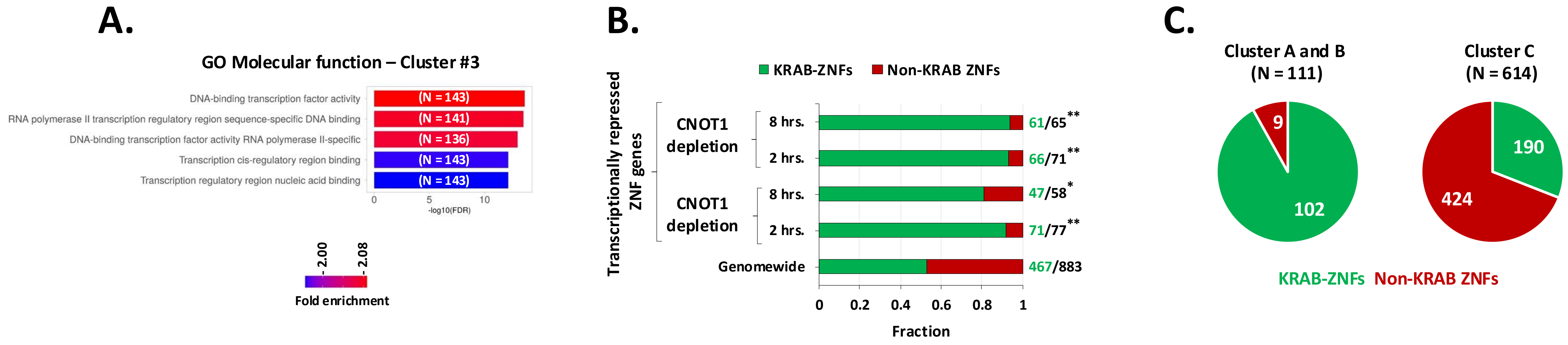

**Supplemental Figure legends**

**Supplemental Figure S1. Genotyping, confirmation of expression, depletion in early timepoints. (A)** Schematic showing the modified genomic loci of CNOT1 and 4. The repair construct depicts left homology arm (LHA), right homology arm (RHA) and sequences coding for auxin inducible degron-T2A peptide-Hygromycin resistance (AID-T2A-HYG-R) cassette. Primers used for genotyping ($P_{out}$ and $P_{in}$) and the stop codon (STOP) in the open reading frame are shown. **(B, C)** Genomic DNA PCR using genomic DNA from parental cells (DLD-1 cells with a TIR1 insertion at the AASV locus, see methods) or cells tagged with an AID tag. CNOT1 clone 7 and CNOT4 clone 3 are heterozygous. **(D)** Trypan blue dye exclusion test to assay cell viability of parental, $CNOT1^{AID}$ or $CNOT4^{AID}$ cells in the absence of auxin. **(E, F)** Tagging CNOT1 or CNOT4 does not affect CCR4-NOT complex subunit accumulation. Western blot analysis of subunits of CCR4-NOT complex in parental and $CNOT1^{AID}$- (**E**) or $CNOT4^{AID}$-cells (**F**) with two different amounts of lysates (differing by a factor of 2). Lysates prepared from two different clones for both $CNOT1^{AID}$ and $CNOT4^{AID}$ cells were analyzed. **(G, H)** Depletion of target protein occurs within 60 minutes of auxin treatment. Western blot analysis of parental and $CNOT1^{AID}$ or $CNOT4^{AID}$ cells using antibodies against CNOT1 (**G**) or CNOT4 (**H**) and tubulin was used as a loading control. Cells were treated with 1 mM auxin for 0, 15, 30 and 60 minutes followed by immunoblotting for the proteins indicated in the panel.

**Supplemental Figure S2. Effects of CNOT1 and CNOT4 depletion on cell growth and the cell cycle. (A)** Growth of parental, $CNOT1^{AID}$ and $CNOT4^{AID}$ cells was measured by MTT assay. 5000 cells were seeded in a 96-well plate. After 24 h the medium was replaced by fresh medium ±auxin. MTT assay was performed at 0, 24, 48 and 72 h (Error bars represent SEM; N = 8). (**B)** Cell cycle analysis of parental, $CNOT1^{AID}$ and $CNOT4^{AID}$ cells. Cells were cultured in ±auxin containing media for 8 and 24 h, stained with propidium iodide and analyzed by flow cytometry using a BD LSR-Fortessa instrument. The panels represent quantification of cells in each phase of cell cycle using FlowJo software (https://www.flowjo.com/). Error bars represent SEM; $N \geq 2$, * indicates $p < 0.05$ determined by unpaired two-tailed Student's t-test.

**Supplemental Figure S3. Effects of CNOT1 and CNOT4 depletion on the levels of CCR4-NOT complex proteins. (A, B)** Western blot analysis of $CNOT1^{AID}$ cells (**A**) or $CNOT4^{AID}$ cells (**B**) using antibodies against CCR4-NOT subunits. Cells were treated in ±auxin containing media for times indicated above the panels, followed by immunoblotting.

**Supplemental Figure S4. Identification of CNOT1 and CNOT4 interacting proteins by proximity labelling (BioID). (A, B)** The BioID tagged proteins are expressed to similar levels as the endogenous proteins. Western blot analysis of parental, $CNOT1^{mTID}$ and $CNOT4^{TID}$ cells. Cells were treated with ±biotin for 1 h. Immunoblotting for CNOT1 (**A**) or CNOT4 (**B**) and tubulin as a loading control. Change in migration of CNOT1 or CNOT4 is caused by the mTID or TID fusion. Asterisk in **B** represents a non-specific band detected by anti CNOT4 antibody. **(C, D)** Volcano-plots of $log_2$Fold-enrichment (FE) in relative protein intensity of biotinylated proteins versus negative $log_{10}$ of $p^{adj}$-values. Datapoints in red represent proteins meeting a ≥ 1.5 enrichment and p-adj < 0.05 cutoff. CCR4-NOT complex subunits are marked with a black circle with CNOT1 and CNOT4 indicated by arrows. Each experimental condition had two biological replicates, which displayed high correlations of peptide abundances (R = 0.99, Data not shown). **(C)** Comparison of proteins from $CNOT1^{mTID}$ versus control cells (untagged). **(D)** Data for the comparison between $CNOT4^{TID}$ versus control cells. **(E)** Gene Ontology terms of 'cellular component' of enriched proteins meeting the cutoff values in **C** ($CNOT1^{mTID}$/control). Proteins dataset was analyzed using ShinyGO 0.80 and GO categories with an enrichment FDR < 0.01 are shown. **(F)** Same as in **E**, except the CNOT4-enriched proteins ($CNOT4^{TID}$/control). **(G)** Venn diagram showing the overlap of CNOT1 and CNOT4 enriched proteins. The p-value for the overlap was calculated using RStudio.

**Supplemental Figure S5. Effects of auxin treatment to parental (untagged cells) and analysis of DEGs upon depletion of CNOT1 and CNOT4. (A, B)** Volcano-plots of gene expression in parental cells. The FCs were calculated (as in Figure 3) for parental cells after treatment in ±auxin containing media for 8 h (**A**) or 24 h (**B**). The numbers represent differentially expressed genes with a FC ≥ 1.5 and p-adj < 0.01 (up-regulated in red and down-regulated in blue). For **A** and **B**, each dot represents an individual RNA. **(C)** Numbers of DEGs for all comparisons in steady-state

RNA-seq experiment. DEGs at 2, 8 or 24 h depletions were analyzed together. **(D)** Distribution of DEGs in steady-state RNA-seq described in Figure 3. The DEGs at 2, 8 and 24 h of depletion of CNOT1 (left panel) or CNOT4 (right panel) were analyzed using an online tool - ShinyGO 0.80. The subclass 'other RNAs' includes minority RNA types e.g. miRNA, processed_pseudogene, scRNA, snoRNA, snRNA as assigned by the tool.

**Supplemental Figure S6. Gene ontology analysis of DEGs under depletion of CNOT1 or CNOT4.** **(A, B)** GO analysis of DEGs in steady-state RNA-seq. The gene sets were analyzed using ShinyGO 0.80 and top 10 categories enriched in GO - biological processes are shown. For **A** and **B** - both up-regulated and down-regulated DEGs were analyzed together. **(A)** DEGs upon CNOT1 depletion. **(B)** DEGs upon CNOT4 depletion. Since there were no significant changes at 8 h, only the 2 h and 24 h are shown. **(C)** Venn diagram showing overlap between DEGs upon CNOT1 and CNOT4 depletion (2 h) with p-value of overlap. **(D)** Same as in **C**, except DEGs upon CNOT1 and CNOT4 depletion (24 h) were analyzed.

**Supplemental Figure S7. Supporting data for TPL based half-life determination (RT-qPCR), correlation with other datasets and foldchange in HLs boxplots.**

**(A)** Pearson's correlation coefficient analysis between RNA half-life values in this study from control cells and those from previously published datasets[1–5]. The p-value of significance of correlation and the number of datapoints in the comparison are shown. **(B)** Boxplot analysis of changes in RNA half-lives by type. RNAs were binned based on their type (as described earlier in Figure **S5D**), such as protein coding mRNAs (N ≈ 11,500), lncRNAs (N ≈ 3,000) and other RNAs (N ≈ 2,000). p-values calculated by the Mann Whitney Test are shown. **(C-E)** RT-qPCR of histone mRNAs. Histone H1.0 is a polyadenylated message. RNA levels were normalized to 18S rRNA and then relative expression (with respect to 0 hr. timepoint, set as 1) was plotted. (Error bars represent average deviation, N = ≥ 2).

**Supplemental Figure S8. RT-qPCR validation of candidate mRNAs**. **(A-I)** Parental, $CNOT1^{AID}$, $CNOT4^{AID}$ cells were treated as in Figure 4A, followed by RT-qPCR analysis for multiple candidate mRNAs. The average values are plotted as in S7C (Error bars represent SEM, N = 3).

**Figure S9. Effects of CNOT1/4 depletion on RNA half-lives are affected by RNA half-lives under steady-state conditions and codon optimality content. (A)** Boxplot of fold-change in RNA half-lives upon depletion of CNOT1. RNAs were binned into quartiles of half-lives calculated under control conditions (N = 15,850) and fold-change in RNA half-lives were plotted. Q1 represents highly unstable RNAs (median half-life under control conditions ~ 3.7 h) and Q4 represents highly stable RNAs (median half-life in under control conditions ~ 45 h). * p-value < $10^{-51}$ by Mann Whitney test (compared to 'All'). **(B)** Same as in **B**, except data for CNOT4 depletion were plotted (total N = 15,850). **(C)** Scatterplot of transcript average codon stability coefficients (CSC score) from versus RNA half-lives calculated under control conditions (parental cells). Only mRNAs represented in both studies were used. **(D)** Scatterplot of transcript average PLS2 (PLS score) from versus RNA half-lives calculated under control conditions (parental cells). For **C** and **D** - Total number of datapoints (N), Spearman Rank correlation coefficient (rho) and 2-sided p-value of significance for correlation are shown. **(E, F)** Boxplot of changes in mRNA half-lives upon CNOT1 depletion. The horizontal dotted line represents no change in half-life. P-values by Mann Whitney Test. **(E)** 'All' represents all mRNAs whose changes in half-lives could be calculated upon CNOT1 and CNOT4 depletion, and whose CSC data are available (N = 1007). 'CSC - lowest' represents decile of mRNAs with lowest CSC scores (N = 100, median CSC = -0.015, median half-life = 5.6 h) and 'CSC - highest' represents decile of mRNAs with highest CSC scores (N = 100, median CSC = +0.011, median half-life = 26.3 h). **(F)** 'All' represents all mRNAs whose changes in half-lives could be calculated upon CNOT1 and CNOT4 depletion, and whose PLS2 data are available (N = 10,296). 'PLS - lowest' represents decile of mRNAs with lowest PLS scores (N = 1029, median PLS = -4.4, median half-life = 6.9 h) and 'PLS - highest' represents decile of mRNAs with highest PLS scores (N = 1029, median PLS = +3.4, median half-life = 25.9). **(G)** Same as in **E**, except changes in mRNA half-lives upon CNOT4 depletion were analyzed. **(H)** Same as in **F**, except changes in mRNA half-lives upon CNOT4 depletion were analyzed.

**Supplemental Figure S10. Depletion of CNOT1 or CNOT4 results in wide-spread increase in nascent transcription. (A)** Schematic depicting the experimental setup to study nascent transcription by transient transcriptome sequencing (TT-seq). **(B)** Boxplot analysis of fold-change (FC) in RNA transcription determined by TT-seq. The FCs were determined by DESeq2 analysis of the spike-in (4-thiouracyl labelled *S. pombe* total RNA) normalized RNA-Seq data (depleted versus non-depleted [auxin-treated parental cells]). Total number of datapoints (N) and median fold change (M) are mentioned for each group. **(C-F)** Volcano-plots of $\log_2$FoldChange (FC) in nascent transcription versus negative $\log_{10}$ of p-values based on DESeq2 analysis mentioned in **B**. The numbers represent differentially expressed genes with a FC ≥ 1.5 and p-adj < 0.01. **(G)** Numbers of DTGs for all comparisons. 'Up' represents DTGs showing increased nascent transcription (FC ≥ 1.5, $p^{adj} < 0.01$) and 'Down' represents DTGs showing decreased nascent transcription (FC ≥ 1.5, $p^{adj} < 0.01$) at 2 or 8 h depletion conditions. **(H, I)** Distribution of DTGs in TT-seq. DTGs (up or down-regulated) at 2 or 8 h of depletion were analyzed together. They were classified based on RNA type (as described earlier in Figure **S5D**). DTGs in CNOT1- (**H**) or CNOT4-depleted cells (**I**).

**Supplemental Figure S11. (A)** Gene ontology analysis of genes in cluster 3 in Figure **5A** (N = 2623). The gene set was analyzed using ShinyGO 0.80 and top 5 GO categories based on enrichment FDR are shown. **(B)** Fraction of transcriptionally repressed ZNFs containing a KRAB domain. P-values were calculated by Fisher exact test (* two-tailed p-value < 0.05, ** two-tailed p-value < 0.005). **(C)** Distribution of ZNF genes by type in the multiple clusters defined by k-means clustering in Figure **5C**.

#### Supplementary references


1. Poetz, F. *et al.* RNF219 attenuates global mRNA decay through inhibition of CCR4-NOT complex-mediated deadenylation. *Nat. Commun.* **12**, 7175 (2021).
2. Narula, A., Ellis, J., Taliaferro, J. M. & Rissland, O. S. Coding regions affect mRNA stability in human cells. *RNA* **25**, 1751–1764 (2019).

3. Gillen, S. L. *et al.* Differential regulation of mRNA fate by the human Ccr4-Not complex is driven by coding sequence composition and mRNA localization. *Genome Biol.* **22**, 284 (2021).
4. Tani, H. *et al.* Genome-wide determination of RNA stability reveals hundreds of short-lived noncoding transcripts in mammals. *Genome Res.* **22**, 947–56 (2012).
5. Arango, D. *et al.* Acetylation of Cytidine in mRNA Promotes Translation Efficiency. *Cell* **175**, 1872-1886.e24 (2018).
